# Functional network analysis identifies multiple virulence and antibiotic resistance systems in *Stenotrophomonas maltophilia*

**DOI:** 10.1101/2023.05.15.540742

**Authors:** Larina Pinto, Rajesh P Shastry, Shivakiran Alva, R. Shyama Prasad Rao, Sudeep D Ghate

## Abstract

*Stenotrophomonas maltophilia*, an emerging multidrug-resistant opportunistic bacterium in humans is of major concern for immunocompromised individuals for causing pneumonia and bloodborne infections. This bacterial pathogen is associated with a considerable fatality/case ratio, with up to 100%, when presented as hemorrhagic fever. It is resistant to commonly used drugs as well as to antibiotic combinations. In-silico based functional network analysis is a key approach to get novel insights into virulence and resistance in pathogenic organisms. This study included the protein-protein interaction (PPI) network analysis of 150 specific genes identified for antibiotic resistance mechanism and virulence pathways. Eight proteins, namely, *pilL*, *fliA*, *Smlt2260*, *Smlt2267*, *cheW*, *Smlt2318*, *cheZ*, and *fliM* were identified as hub proteins. Further docking studies of selected phytochemicals were performed against the identified hub proteins. Deoxytubulosine and Corosolic acid were found to be potent inhibitors of hub proteins of pathogenic *S. maltophilia* based on protein-ligand interactive study. Further pharmacophore studies are warranted with these molecules to develop them as novel antibiotics against *S. maltophilia*.

## 1. INTRODUCTION

*Stenotrophomonas maltophilia* is a waterborne aerobic Gram-negative bacterium that is rod-shaped and motile due to polar flagella. *S. maltophilia* has known to be an emerging pathogenic in immunocompromised people (Patterson et al., 2020). Exposure to this bacterium can happen both within and outside of the clinical environment (Brooke, 2012). The two most typical manifestations, bacteremia and pneumonia, have both been linked to considerable death rates during the past 20 years (Senol, 2004). Other clinical syndromes associated with this bacterium are skin and soft-tissue infections (Sakhnini et al., 2002), endocarditis, urinary tract infection, meningitis, mastoiditis, etc. (Senol, 2004).

According to estimates, there are between 5.7 and 37.7 infections for every 10,000 hospital discharges worldwide, which is considerably greater than previously thought during the years since the 1970s (Patterson et al., 2020; Said et al., 2022). The rise in immunocompromised individuals and widespread usage of broad-spectrum antibiotics are believed to be the principal causes for this growing infection rate (Said et al., 2022).

Infection management efforts are made more difficult by *S. maltophilia*’s capacity to grow biofilms on biotic surfaces and fomites. Additionally, the blurred lines between colonization and infection, and the frequent polymicrobial presentation of *S. maltophilia*, particularly in immunocompromised hosts, lead to delay in the administration of the proper antimicrobial therapy. In turn, this contributes to the overuse and abuse of antibiotics in cases of non-infection without appropriate diagnosis. Furthermore, the abundance of innate and acquired resistance mechanisms restrict the range of curative alternatives (Kullar et al., 2022).

*Stenotrophomonas* is intrinsically resistant to an assortment of antibiotics, including carbapenems, aminoglycosides, macrolides, β-lactams, tetracyclines, trimethoprim-sulfamethoxazole (TMP-SMX), chloramphenicol, and fluoroquinolones (Appaneal et al., 2020). Some of the most important molecular variables affecting this organism’s resistance to antibiotics are the expression of qnr genes, the generation of β-lactamases, the presence of class 1 integrons, and efflux pumps. While most studies show that *S. maltophilia* is sensitive to TMP/SMX (Chang et al., 2015), a few studies have found resistance indicating the emergence of antimicrobial resistance (AMR) in *S. maltophilia* (Patterson et al., 2020; Saleh et al., 2021).

Given the emergence of AMR in *S. maltophilia*, drug resistance determinants are of great interest. Crossman et al. (2008) found nine potential antimicrobial efflux systems of the resistance-nodulation-division (RND) type were present, in addition to a number of genes that confer resistance to antimicrobial drugs of different sorts via other pathways.

In this work, the opportunistic pathogen *S. maltophilia K279a* was investigated using gene interaction network analysis to look into several antimicrobial resistance (AMR) and virulence genes. The exceptional ability of this specific strain to withstand drugs and heavy metals along with its pathogenicity were the reasons why it was chosen. We found biologically relevant genes involved in resistance and virulence mechanisms. Prospectively, this research will benefit wet lab researchers in designing cutting-edge therapeutic approaches to counteract *S. maltophilia* pathogenicity.

## 2. METHODOLOGY

### 2.1 Sequence acquisition

The complete genome reference sequences (RefSeqs, a total of 72) for all *S. maltophilia* were downloaded from the NCBI website (https://www.ncbi.nlm.nih.gov/, last accessed on Apr 27, 2023). Further, sequence related metadata including the geographical origins of the isolates were also collected from the NCBI.

### 2.2 Identification of virulence and antibiotic resistance genes

Each genome sequence was submitted to the resistance gene identifier (RGI) tool of comprehensive antibiotic resistance database (CARD) (Alcock et al., 2020) to obtain annotations based on perfect, strict, or loose paradigm, and complete gene match criteria for the identification of antibiotic resistance genes (Rao et al., 2023). The virulence factor database (VFDB, http://www.mgc.ac.cn/VFs/) is a comprehensive warehouse and online platform widely used for the identification of virulence factors (VFs) (Liu et al., 2019). We used Abricate 0.9.8 (https://github.com/tseemann/abricate) interface to screen the VFDB using the parameters – percentage identity of ≥60[%[and coverage of ≥40[%. The resulting lists of genes were combined and duplicate entries were removed. They were then validated by comparing against the protein-coding genes from the whole genome of *S. maltophilia K279a* strain (NC_010943.1).

### 2.3 Construction of protein-protein interaction (PPI) network

In order to investigate the interacting links between the AMR and virulent genes in *S. maltophilia K279a* strain, they were mapped using Search Tool for the Retrieval of Interacting Genes database (STRING v.11.0, https://string-db.org). Visualization of the PPI network was carried out using Cytoscape (https://cytoscape.org). The PPI network was constructed with a high confidence level of 0.7 and then imported to Cytoscape (3.9.2) for further analysis.

### 2.4 Topological analysis

The PPI network was analyzed for its topological properties using Network Analyzer, a Cytoscape plugin. This tool analyses all the aspects of the PPI network. To better understand the interaction between the proteins, connectivity degree (K), betweenness centrality (BC) and closeness centrality (CC) were analyzed. These parameters provide the information about the number of proteins connected to each protein, centrality of the proteins, and the distance between them.

### 2.5 Functional and pathway enrichment analysis of AMR and virulence proteins

ClueGO, a Cytoscape plug-in, was used for thorough analysis and visualization of the functionally enriched set of proteins. With a p-value of ≤0.05, STRING was used to obtain annotations and Gene Ontology (GO) concepts for genes and their functional relationships. KEGG (Kanehisa and Goto, 2000), UniProt, Pfam, and InterPro were used to comprehend critical pathway information of the genes and proteins involved in diverse activities as described in earlier study (Shetty et al., 2022).

### 2.6 Screening of hub genes and clusters

The PPI network was screened for its hub genes using a Cytoscape plug-in, CytoHubba (Chin et al., 2014). Top 20 genes were identified from the PPI network using each of 12 in-built algorithms of CytoHubba. The genes that overlapped in more than 6 algorithms were identified as the hub genes in the PPI network and were assumed to play a critical role in AMR and virulence. A Cytoscape plugin, Molecular Complex Detection (MCODE), with parameters set as the degree threshold (2), node score threshold (0.2), k-core threshold (4-6), and max depth of network (100) with other default parameters, was used to screen for deeply linked clusters within the PPI network. Based on the results, a suitable k-core was selected for further analysis. A subnetwork was generated with the selected nodes of the clusters from MCODE results including all edges along with seed proteins. These hub genes/proteins were considered as potential druggable targets/models, and taken for model quality assessment/evaluation and docking.

### 2.7. Model quality assessment and evaluation

Model validation servers were utilized to analyze the physicochemical features of the identified hub proteins and determine metrics such as Z-score, Q mean DisCo Global score, Ramachandran scores, etc. The functions of hub proteins were validated and their cellular localization was predicted to understand their function as per the methodology described earlier (Shetty et al., 2022).

### 2.8. Docking studies

The phytochemically derived molecules that might act as inhibitors of hub proteins based on the information that those molecules were used in the treatment of respiratory infections were selected from IMPPAT library. Docking studies with selected molecules and hub proteins were conducted to understand the chemistry of interaction. The structure of hub proteins was obtained from AlphaFold. PyRx virtual screen tool version 0.9.8 (Dallakyan and Olson, 2015) was used for the docking of the phytochemical inhibitors of hub proteins. The ligands (selected inhibitors) were retrieved from PubChem and used to create the 3D structure (Shetty et al., 2022). The ligand energy was then minimized, and a ligand file was created in accordance with the specifications. The software’s requirements were followed for maintaining the docking parameter, and the optimal postures were chosen based on the binding energy. Discovery Studio 2020 Client and UCSF Chimera version 1.10.1 were used to evaluate the output files.

## 3. RESULTS

### 3.1 Reconstruction of AMR and virulence PPI network of S. maltophilia

CARD yielded 75 AMR genes while VFDB yielded 92 virulence genes (Table S1 and S2). We selected the *S. maltophilia K279a* strain because of it being the core *S. maltophilia* genome in STRING. All virulence genes were present, but only a small subset of the AMR genes (28 genes) was available in the STRING database. The network was extended by setting the total number of interactors to 50. The final network consisted of 150 genes with 1479 functional interactions.

After removing the loosely bound connections, the PPI network was visualized and analyzed with Network Analyzer (Figure 1A and 1B). MCODE was used to screen the PPI network for highly interconnected clusters, which resulted in the PPI network being divided into 3 clusters viz. C1 (40 nodes, 745 edges, seed protein *motA*), C2 (8 nodes, 26 edges, seed protein *Smlt2823*), C3 (16 nodes, 49 edges, seed protein *rpoA* (Figure 2). By using 12 distinct CytoHubba metrics, proteins that overlapped in 7 or more parameters were classified as top hub proteins and eight such hub proteins (*pilL, fliA, Smlt2260, Smlt2267, cheW, Smlt2318, cheZ,* and *fliM*) were identified (Table S2). These hub proteins were selected for docking analysis.

**Figure 1:**
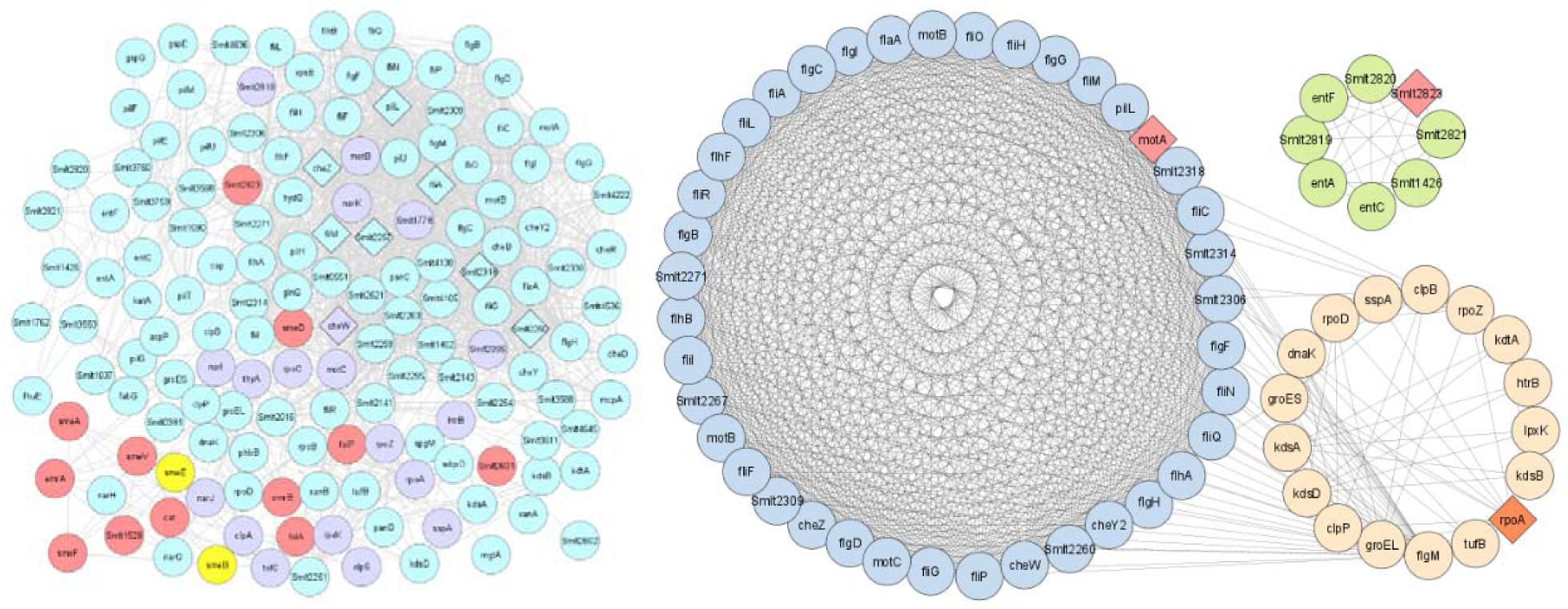
(A) Protein-protein interaction network between virulence (blue-colored) and AMR (pink-colored) nodes/proteins. Proteins categorized as both virulence and AMR are colored in yellow while extended proteins are colored in purple. Rhomboids are used to represent each of the eight hub proteins. The network has 150 nodes connected by 1479 edges. (B) Three clusters were identified (using MCODE) from the protein-protein interaction network with a MCODE score of >6. Cluster 1 (blue-colored nodes; seed protein motA, score: 32.6), Cluster 2 (green-colored nodes, seed protein Smlt2823, score: 6.0) and Cluster 3 (orange-colored nodes, seed protein rpoA, score: 5.1). Seed protein is represented in pink-colored rhomboid.

**Figure 2:**
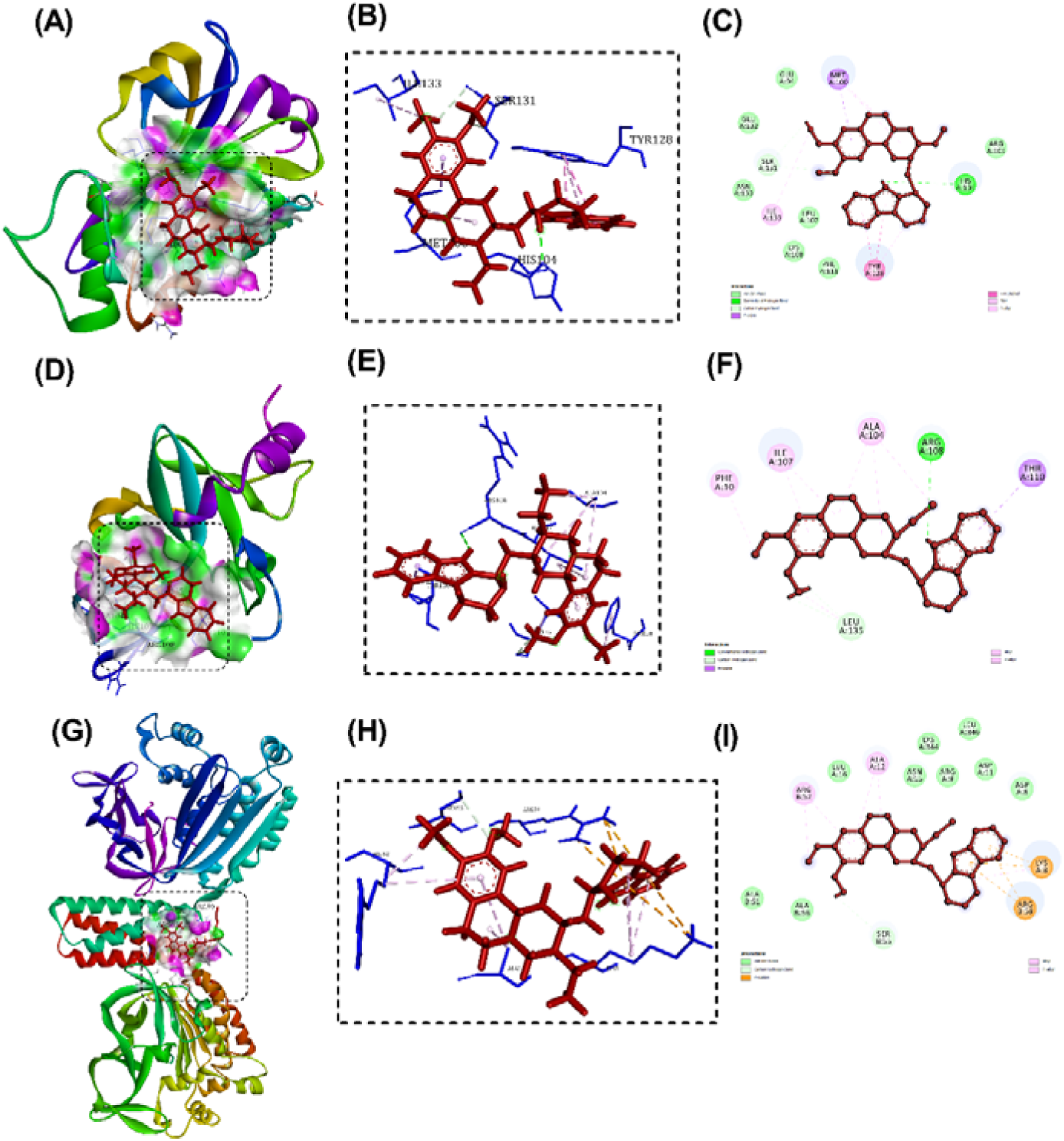
Molecular docking of FliM, CheW, and Smlt2267 with Deoxytubulosine. Binding confirmation of proteins (A, D, and G), snapshot of ligand-protein complexes (B, E, and H), and 2D interactions of ligand with respective amino acids (C, F, and I) are shown.

**Figure 3:**
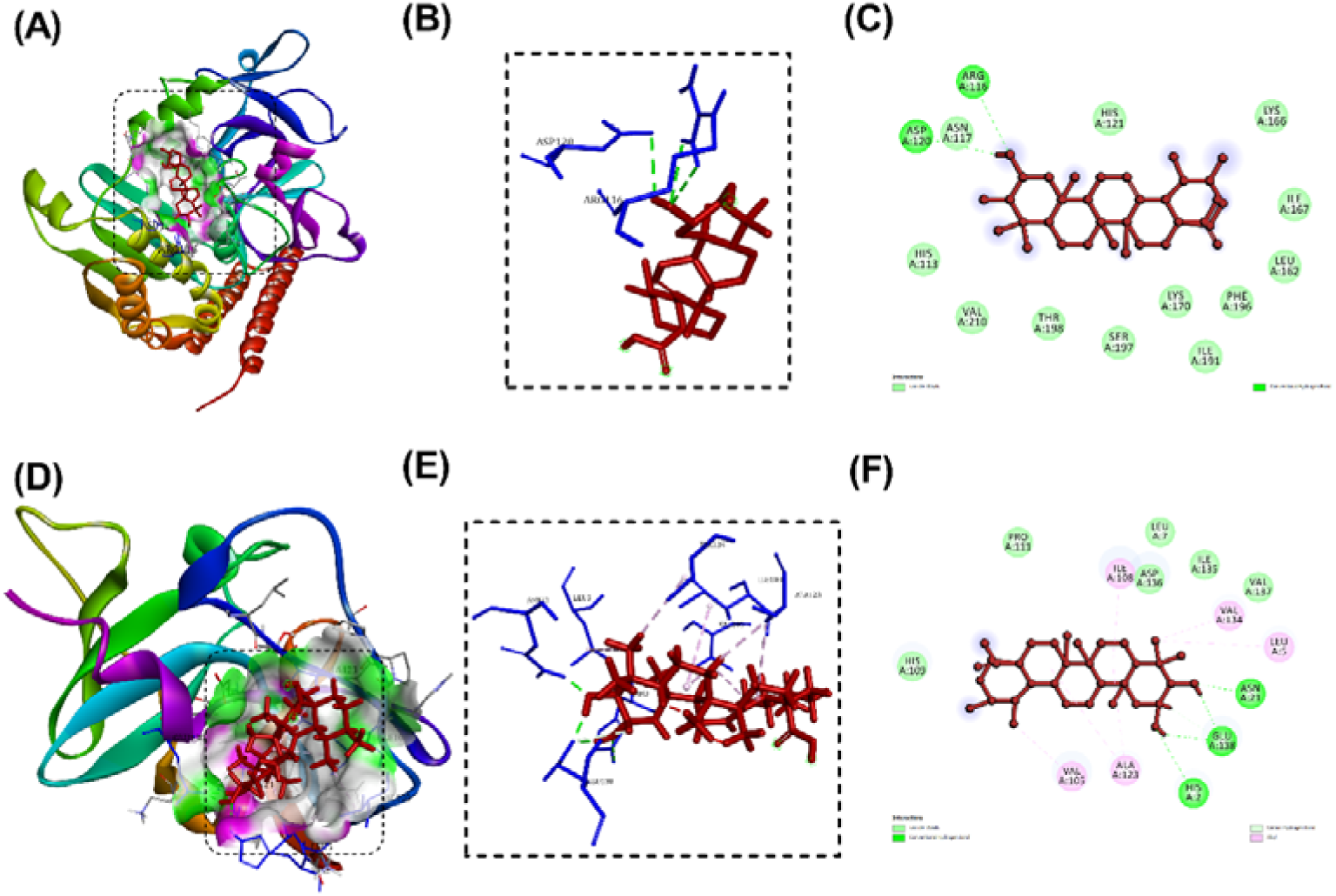
Molecular docking of Smlt2260 and Smlt2318 with Corosolic acid. Binding confirmation of proteins (A and D), snapshot of ligand-protein complexes (B and E), and 2D interactions of ligand with respective amino acids (C and F) are shown.

**Figure 4:**
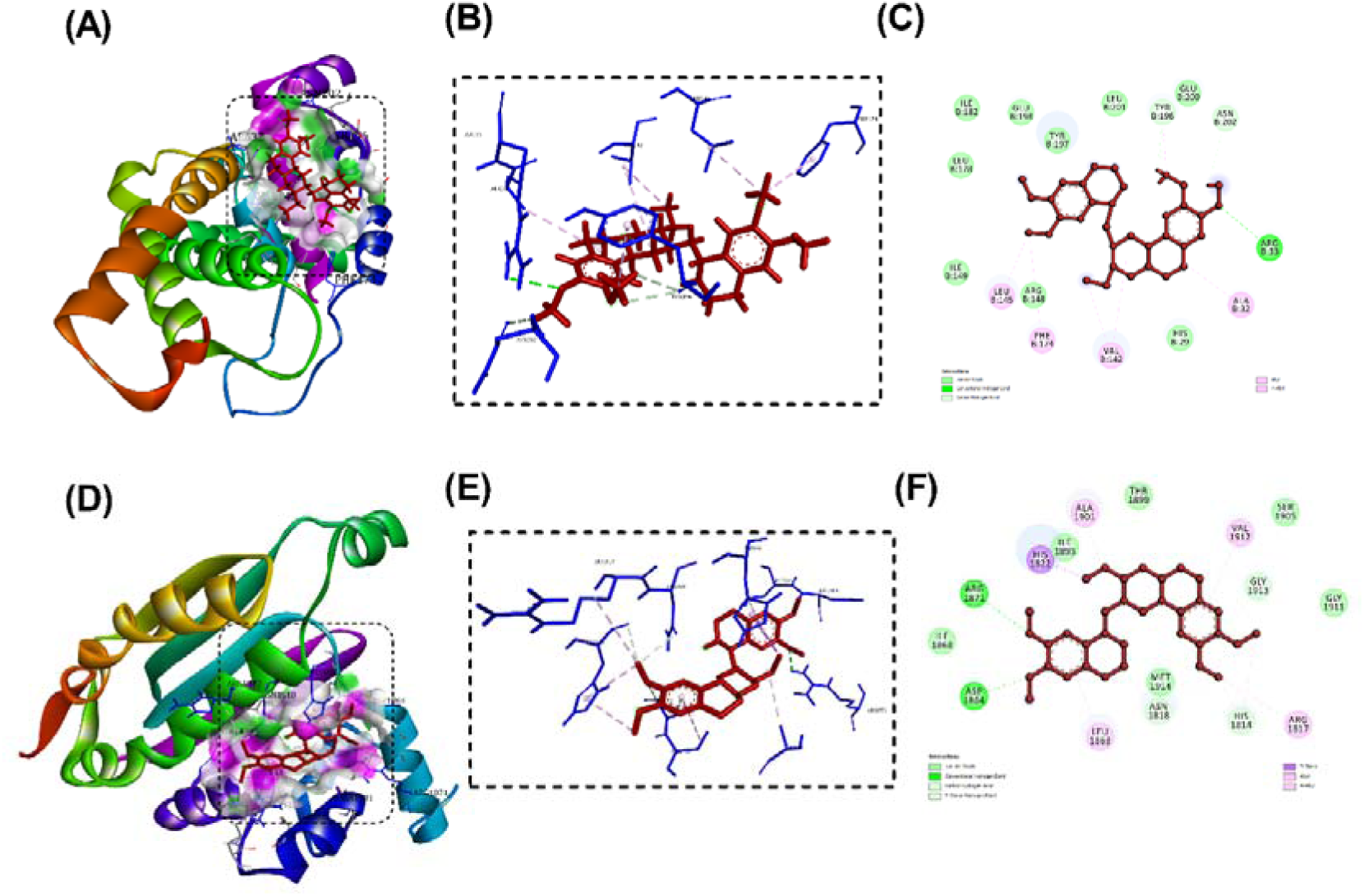
Molecular docking of FliA and PilL with Emetine. Binding confirmation of proteins (A and D), snapshot of ligand-protein complexes (B and E), and 2D interactions of ligand with respective amino acids (C and F) are shown.

### 3.2 Topological features of PPI network

The protein interaction network (PIN) can be assessed by its mutual connections and topology. The topology of the full network and subnetwork were analyzed using Network Analyzer (Table 1). The BC of the full network and subnetwork were found to be 0.0135 and 0.01857, while the CC were 0.3493 and 0.57928, respectively. The analysis revealed that the hub protein *pilL* had the highest degree value and CC value, while *Smlt2141* had high BC value (Table 2). Clustering coefficient represents the closeness of nodes and neighbors, and the hierarchical modularity of the PIN, and is used to spot the possible functional modules and uncover the molecular complexes or signaling pathways in the PIN. The clustering coefficients were 0.8097 and 0.549 for the full network and subnetwork, respectively. The average number of neighbors for 150 nodes was 19.7.

**Table 1.** MCODE parameters and topological parameters of whole network, clustered genes, and individual clusters using Network Analyzer.

| Parameters | Network |  | Clusters |  |  |
| --- | --- | --- | --- | --- | --- |
|  | Whole network | Subnetwork | C1 | C2 | C3 |
| Number of Nodes | 150 | 64 | 40 | 8 | 16 |
| Number of Edges | 1479 | 829 | 745 | 26 | 49 |
| Network Density | 12.58 | 0.52 | 0.95 | 0.92 | 0.40 |
| Clustering Coefficient | 0.66 | 0.9 | 0.96 | 0.94 | 0.84 |
| Average Number of Neighbors | 19.72 | 28.67 | 37.25 | 6.5 | 6.12 |
| Characteristic path length | 2.99 | 2.05 | 1.04 | 1.07 | 2.01 |
| Network Diameter | 8 | 5 | 2 | 2 | 4 |
| Betweenness centrality | 0.0135 | 0.0185 | - | - | - |
| Closeness centrality | 0.3493 | 0.5792 | - | - | - |
| Genes present in clusters |  |  | fliQ | Smlt2823 | rpoD |
|  |  |  | CheW | Smlt2820 | rpoA |
|  |  |  | motA | Smlt1426 | sspA |
|  |  |  | flhB | entF | clpP |
|  |  |  | flaA | entA | kdsB |
|  |  |  | fliL | Smlt2819 | tufB |
|  |  |  | Smlt2318 | entC | kdsD |
|  |  |  | motB | Smlt2821 | groEL |
|  |  |  | fliM |  | lpxK |
|  |  |  | Smlt2306 |  | groES |
|  |  |  | fliO |  | kdsA |
|  |  |  | cheZ |  | rpoZ |
|  |  |  | flhA |  | htrB |
|  |  |  | fliC |  | clpB |
|  |  |  | cheY2 |  | dnaK |
|  |  |  | flgC |  | kdtA |
|  |  |  | Smlt2309 |  |  |
|  |  |  | fliI |  |  |
|  |  |  | Smlt2314 |  |  |
|  |  |  | flgH |  |  |
|  |  |  | flgI |  |  |
|  |  |  | fliN |  |  |
|  |  |  | fliR |  |  |
|  |  |  | flgF |  |  |
|  |  |  | fliF |  |  |
|  |  |  | MotA |  |  |
|  |  |  | flgD |  |  |
|  |  |  | fliP |  |  |
|  |  |  | MotB |  |  |
|  |  |  | fliG |  |  |
|  |  |  | fliH |  |  |
|  |  |  | pilL |  |  |
|  |  |  | flhF |  |  |
|  |  |  | flgG |  |  |
|  |  |  | cheA |  |  |
|  |  |  | flgM |  |  |
|  |  |  | CheA |  |  |
|  |  |  | flgB |  |  |
|  |  |  | fliA |  |  |
|  |  |  | Smlt2271 |  |  |

**Table 2.**
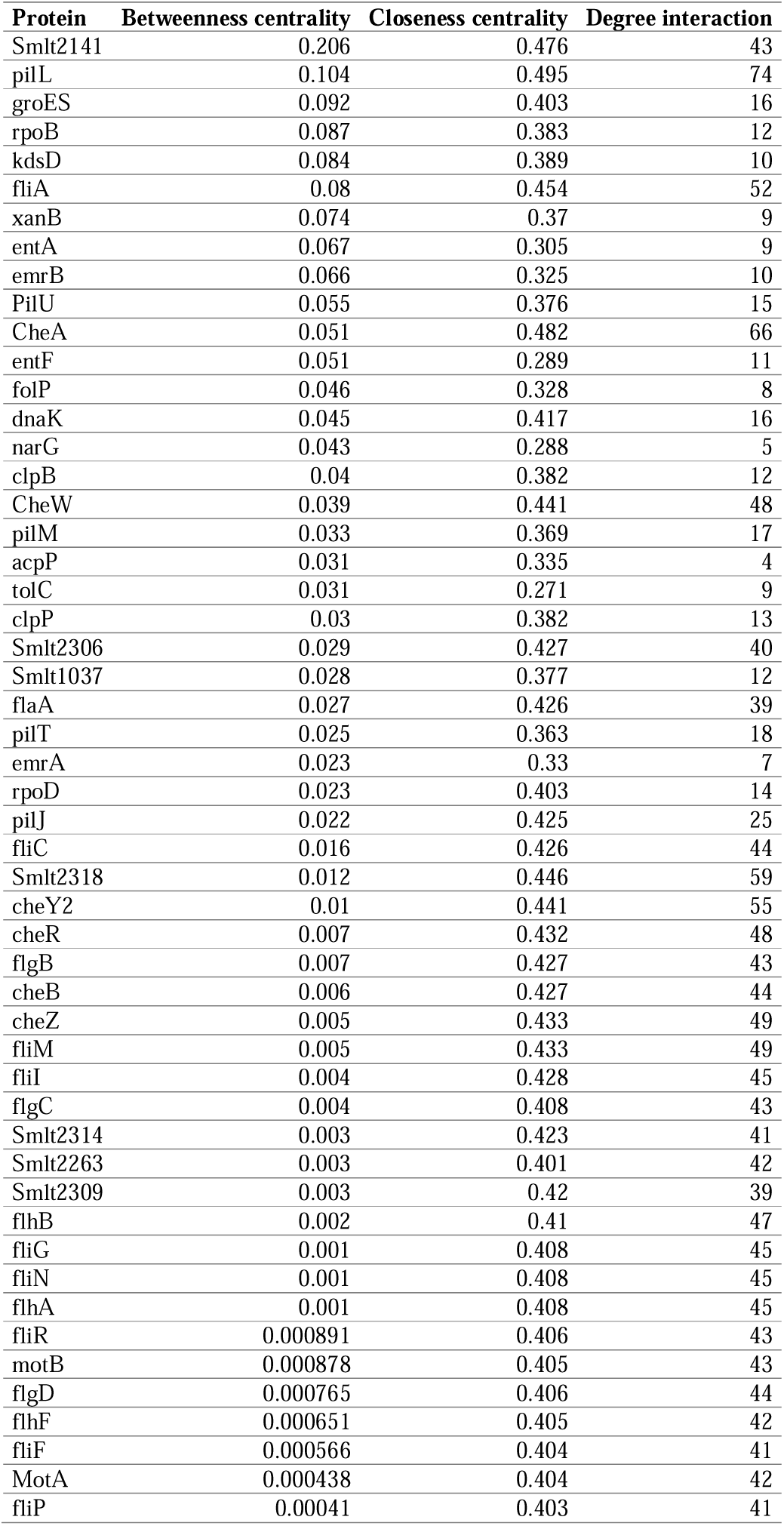
List of proteins with highest betweenness centrality, closeness centrality, and degree interaction.

| Protein | Betweenness centrality | Closeness centrality | Degree interaction |
| --- | --- | --- | --- |
| Smlt2141 | 0.206 | 0.476 | 43 |
| pilL | 0.104 | 0.495 | 74 |
| groES | 0.092 | 0.403 | 16 |
| rpoB | 0.087 | 0.383 | 12 |
| kdsD | 0.084 | 0.389 | 10 |
| fliA | 0.08 | 0.454 | 52 |
| xanB | 0.074 | 0.37 | 9 |
| entA | 0.067 | 0.305 | 9 |
| emrB | 0.066 | 0.325 | 10 |
| PilU | 0.055 | 0.376 | 15 |
| CheA | 0.051 | 0.482 | 66 |
| entF | 0.051 | 0.289 | 11 |
| folP | 0.046 | 0.328 | 8 |
| dnaK | 0.045 | 0.417 | 16 |
| narG | 0.043 | 0.288 | 5 |
| clpB | 0.04 | 0.382 | 12 |
| CheW | 0.039 | 0.441 | 48 |
| pilM | 0.033 | 0.369 | 17 |
| acpP | 0.031 | 0.335 | 4 |
| tolC | 0.031 | 0.271 | 9 |
| clpP | 0.03 | 0.382 | 13 |
| Smlt2306 | 0.029 | 0.427 | 40 |
| Smlt1037 | 0.028 | 0.377 | 12 |
| flaA | 0.027 | 0.426 | 39 |
| pilT | 0.025 | 0.363 | 18 |
| emrA | 0.023 | 0.33 | 7 |
| rpoD | 0.023 | 0.403 | 14 |
| pilJ | 0.022 | 0.425 | 25 |
| fliC | 0.016 | 0.426 | 44 |
| Smlt2318 | 0.012 | 0.446 | 59 |
| cheY2 | 0.01 | 0.441 | 55 |
| cheR | 0.007 | 0.432 | 48 |
| flgB | 0.007 | 0.427 | 43 |
| cheB | 0.006 | 0.427 | 44 |
| cheZ | 0.005 | 0.433 | 49 |
| fliM | 0.005 | 0.433 | 49 |
| fliI | 0.004 | 0.428 | 45 |
| flgC | 0.004 | 0.408 | 43 |
| Smlt2314 | 0.003 | 0.423 | 41 |
| Smlt2263 | 0.003 | 0.401 | 42 |
| Smlt2309 | 0.003 | 0.42 | 39 |
| flhB | 0.002 | 0.41 | 47 |
| fliG | 0.001 | 0.408 | 45 |
| fliN | 0.001 | 0.408 | 45 |
| flhA | 0.001 | 0.408 | 45 |
| fliR | 0.000891 | 0.406 | 43 |
| motB | 0.000878 | 0.405 | 43 |
| flgD | 0.000765 | 0.406 | 44 |
| flhF | 0.000651 | 0.405 | 42 |
| fliF | 0.000566 | 0.404 | 41 |
| MotA | 0.000438 | 0.404 | 42 |
| fliP | 0.00041 | 0.403 | 41 |

### 3.3 Functional enrichment analysis

Annotations and Gene Ontology (GO) terms were retrieved for the genes and their functional partners from STRING database with a p-value of ≦ 0.05, where the functional enrichment analysis data of these genes were given. The data provided various properties and functions of AMR as well as virulence genes in the network. The enriched data were from KEGG pathways, Pfam protein domains, UniProt, and InterPro databases. A total of 44 GO terms were collected out of which 27, 9, and 8 terms corresponded to Biological Processes (BP), Molecular Functions (MF) and Cellular Components (CC) respectively. The top biological processes based on the number of genes associated involve cellular anatomical entity [GO:0110165], nucleotide binding [GO:0000166], purine ribonucleotide binding [GO:0032555] and cellular process [GO:0009987].

### 3.4 Model evaluation

ProtParam Tool was implemented to examine the physicochemical characteristics *of pilL, fliA, Smlt2260, Smlt2267, cheW, Smlt2318, cheZ*, and *fliM* proteins (Table 3). *CheW* showed highest aliphatic index value indicating high thermostability. The most unstable was *fliA*, which had the greatest value of the instability index, whereas *pilL* had the highest extinction coefficient. The model evaluation score analysis revealed that except *pilL*, all other proteins carried a negative overall G-factor score. In *pilL, fliA, cheZ*, and *fliM*, the proportion of generously allowed regions was zero, whereas the others had values greater than zero. But the percentage of all generously allowed regions was less than 1%. The protein *Smlt2260* had a better overall quality compared to other hub proteins, as predicted by ERRAT. The Levitt-Gerstein (LG) and MaxSub scores established by ProQ, as well as the resolution estimated by ResProx, demonstrate the dependability of constructed 3D models. The Ramachandran favored percentages of the core hub proteins *pilL, fliA, Smlt2260, Smlt2267* and *fliM* were above 90% (Table 4) suggesting that the protein structures were of great stereochemical quality, as predicted by MOLPROBITY. The Ramachandran plot for *Smlt2260* (protein having highest percentage of favored regions) is given in Fig S1. BLASTp, MOTIF, STRING, and ScanProsite were performed to evaluate the precision of the functions annotated by GenBank. Table S7 lists the annotated functions. All hub proteins were anticipated to be localized to the cytoplasm, according to PSORTb v 3.0.3 and PSLPred, although *fliM* might have several localization sites (Table S7).

**Table 3.** Physicochemical properties of the hub proteins.

| Parameters | **pilL** | **fliA** | **Smlt2260** | **Smlt2267** | **CheW** | **Smlt2318** | **cheZ** | **fliM** |
| --- | --- | --- | --- | --- | --- | --- | --- | --- |
| Theoretical pI | 4.27 | 5.5 | 4.97 | 5.16 | 4.39 | 5.38 | 4.77 | 5.08 |
| Molecular weight (kD) | 237570.7 | 27238.8 | 70281.2 | 65129.4 | 17561.9 | 34244.2 | 21938.6 | 37673.1 |
| Extinction coefficient | 110825* | 14440 | 30940 | 20065* | 5960 | 8940 | 7115* | 27055* |
| Instability index | 44.49<br>(Unstable) | 47.38<br>(Unstable) | 40.37<br>(Unstable) | 41.32<br>(Unstable) | 22.24<br>(Stable) | 44.66<br>(Unstable) | 53.35<br>(Unstable) | 43.34<br>(Unstable) |
| Aliphatic index | 94.75 | 93.72 | 105.84 | 103.25 | 112.45 | 109.87 | 99.1 | 98.89 |
| No. of amino acids | 2225 | 247 | 663 | 609 | 163 | 314 | 201 | 334 |
| Grand average of hydropathicity (GRAVY) | -0.176 | -0.344 | 0.054 | -0.044 | 0.19 | 0.07 | -0.374 | -0.149 |

\*Assuming all pairs of Cys residues form cystines.

**Table 4.**
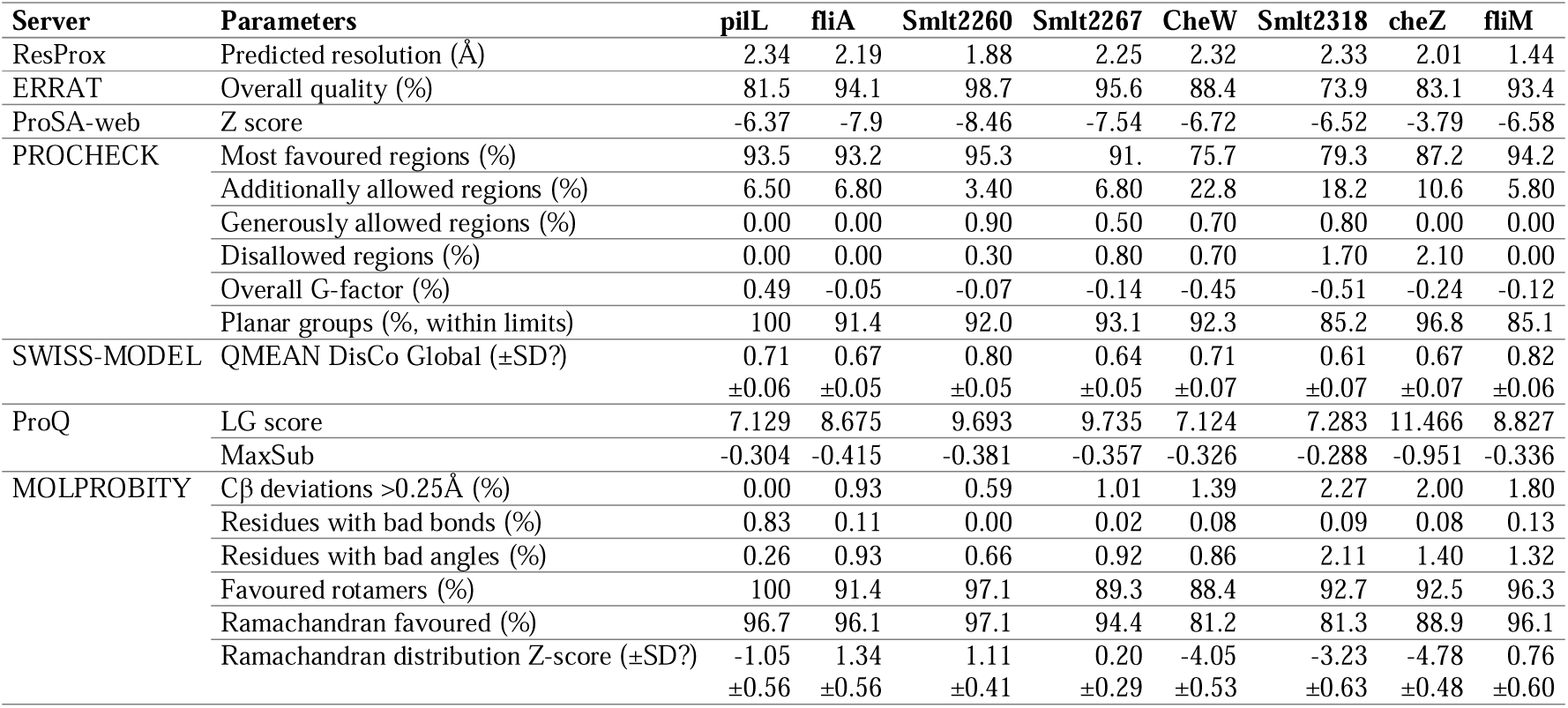
Model evaluation scores of the hub proteins.

### 3.5 Docking analysis

Docking analysis with selected ligands (Table S8) were performed to hub proteins *pilL, fliA, Smlt2260 (cheA), Smlt2267 (cheA2), cheW, Smlt2318* (two-component response regulator chemotaxis signal), *cheZ,* and *fliM* (Table S9). The details of hydrogen bonds and the resulting binding energies for selected chosen ligands are given in Table 5. Out of multiple selected compound Deoxytubulosine showed a lower binding energies with *Smlt2267, cheW*, and *fliM*.

**Table 5.** Molecular binding affinity of selected ligands against hub proteins of *S. maltophilia*.

| Protein name | Ligand | Binding affinity (kcal/mol) | Hydrogen bonds |
| --- | --- | --- | --- |
| CheW | Deoxytubulosine | -8.793 | ARG108 |
| CheZ | Cephaeline | -8.137 | GLN101, ASP102 |
| FliA | Emetine | -8.796 | ARG33 |
| FliM | Deoxytubulosine | -9.579 | HIS104 |
| PilL | Emetine | -7.988 | ASP1864, ARG1871 |
| Smlt2260 | Corosolic acid | -9.174 | ARG393, ASP397 |
| Smlt2267 | Deoxytubulosine | -8.648 | GLN58, ASP148 |
| Smlt2318 | Corosolic acid | -8.833 | HIS2, ASN21, GLU138 |

Furthermore, Corosolic acid favored to bind Smlt2318 and Smlt2260, followed by Emetine to *fliA* and *pilL* proteins of *S. maltophilia*. These effective interactions of selected phytochemical-based ligands to hub proteins suggested a role in structure-activity relationship. The optimal interactions with the lowest autodock score and the best conformation are given in Figures 2, 3, and 4. Based on protein-ligand interactive study, Deoxytubulosine and Corosolic acid might be best candidate inhibitors of hub proteins of pathogenic *S. maltophilia*.

## 4. DISCUSSION

The continued development of antibiotics is an immediate need if humankind is to stay ahead and counter the emergence and spread of antibiotic resistance. As per recent estimates, about 0.7 million deaths annually worldwide are attributed to AMR; with the number projected to grow rapidly to a tune of 10 million deaths per year by 2050 (O’Neill, 2016). Hospital infections present the most pressing need for novel therapeutics. Aside from the current AMR bacterial species, a small but increasing number of isolates, predominantly Gram-negative bacteria (such as *S. maltophilia*), are becoming resistant to previously effective antibiotics available in the market, further exacerbating the issue of antibiotic resistance (Livermore, 2004). Additionally, this bacterium has recently been identified as the most prevalent Gram-negative carbapenem-resistant pathogen isolated from clinical settings. This remains one of the relatively understudied bacteria in comparison to other Gram-negative bacteria despite having an undeniable clinical impact (Cai et al., 2020). Due to *S. maltophilia*’s inherent resistance to several antibiotics and its propensity to acquire additional resistance through horizontal gene transfer and mutation, treatment of this organism can be challenging. The strain’s resistance to quinolones, cotrimoxale, and/or cephalosporins, the antibiotics routinely used to treat *S. maltophilia* infections, have evolved in recent years (Sánchez, 2015).

In order to maintain a steady flow of new antibacterial drug candidates into the development pipeline, it is pivotal to accelerate antibiotic optimization efforts. For this reason, it is necessary to boost the early stages of drug discovery and development since they are crucial for identifying and validating novel therapeutic candidates that can effectively combat antibacterial resistance. The attrition rate in antibacterial drug discovery has been particularly high in recent decades, as evidenced by the fact that no new class of Gram-negative antibiotics has been introduced in more than half-century (Miethke et al., 2021). However, designing entirely new scaffolds is much more expensive than developing derivatives of established compound classes (Schlander et al., 2021). Phytopharmaceuticals, which have recently attracted global interest, can be used to solve the dearth of novel medications in development (Konwar et al., 2022). Antibiotics and plant extracts work together synergistically to fight resistant bacteria, opening up new options for the treatment of infectious disorders. This feature makes it possible to continue using the specific antibiotic even after it loses its therapeutic impact (Sibanda and Okoh, 2007).

The sequenced complete genome of *S. maltophilia* K279a was analyzed in our study. This specific strain sheds information on the potential genetic underpinnings of adaptation to various habitats, which eventually resulted in enhanced host pathogenicity and resistance to a spectrum of drugs (Abda et al., 2015). Understanding bacterial pathogenicity and their interactions with the host, which may also serve as novel targets in pharmaceutical and vaccine development, requires the discovery of virulence factors. Over the past few decades, the advent of post-genomic methods, like genomics, transcriptomics, and proteomics, has sped up the discovery of virulence factors (Mason et al., 2018). We identified prospective pharmacological targets to aid in the development of innovative treatments to address the resistance mechanism.

By generating interaction networks, analyzing clusters, and investigating functional enrichment, this work revealed important information on efflux pumps and biofilm formation, as well as other drug resistance and pathogenicity mechanisms of the *S. maltophilia* K279a strain. The virulence and AMR genes found in this work have been reported previously (Huang et al., 2017). For example, the bacterial outer membrane lipoprotein *pilL* has been linked to pilus production, motility, and genetic transformation in earlier investigations (Sakai et al., 2000). *FliM* is a flagellar protein with diverse roles, while *<u>fliA</u>* has been demonstrated to be a sigma factor specific for class 3 flagellar operons (Eichelberg et al., 2000).

Chemotaxis is one of the known mechanisms that helps bacteria adhere to surfaces and develop by producing biofilms. This has been seen in many habitats and culture settings in various bacteria. Chemotaxis pathways would control both excitation and adaptability to environmental cues since it may be necessary for bacterial survival, metabolism, and interactions within ecological niches. *CheA* (*Smlt2260* and *Smlt2267*) is a critical gene in controlling the onset of bacterial chemotaxis. *CheA* is a methyl-accepting protein that can recognize cues from the environment (Albornoz et al., 2017). Another notable hub protein responsible for chemotaxis in our study is *cheW*, which will help us understand the genetic and biochemical makeup that will be pertinent in the search for new antibiotics (Liu et al., 1991). Another chemotactic protein, *Smlt2318*, possesses operon-like characteristics and stimulates their transcription, which may be a crucial regulatory step in the development of *S. maltophilia* biofilms (Kang et al., 2015). The hub proteins play a crucial role in the PPI network’s operations. They are also implicated in a number of virulence mechanisms and may be a important source of prospective therapeutic targets (Wang et al., 2011). For experimental biologists, the network and sub-networks captured by this topological analytic technique will provide fresh perspectives on crucial regulatory networks and protein drug targets.

Both processes i.e., antimicrobial resistance and virulence mechanisms are traditionally thought to be essential for bacteria to survive in challenging environments from a biological standpoint (Christaki et al., 2020). The development of antimicrobial resistance is crucial for pathogenic bacteria to be able to withstand antimicrobial therapies, overcome host defense mechanisms, and adapt to and flourish in challenging conditions. Virulence mechanisms are required to counter host defense mechanisms (Beceiro et al., 2013). Hub protein structural insights can help in the absence of phenotypic data and can also give a physical foundation for a more thorough understanding of therapeutic targets to tackle antimicrobial resistance (Shetty et al., 2022). As a result, computational methods can help with the mechanistic understanding of how different phytochemicals interact with proteins. Predicting suitable non-traditional compounds can be done by further extending our understanding of the impacts of protein-ligand stability using molecular docking experiments.

The eight hub proteins were created in this study as high-quality, energy-minimized 3D models using SWISS MODEL. On model validation servers, additional parameters and physicochemical characteristics were evaluated. The *Smlt2267* (*CheA*) protein in *Vibrio harveyi* regulate bacterial motility, and adhesion at different temperature and salinity as well as pH values. The role of *RecA* and a *CheW*-like proteins are proved to be required for surface-associated motility as well as virulence of the multi-drug resistant pathogen *Acinetobacter baumannii*. Currently, Antibiotic resistance breakers (ARBs), such as a drug combination, are being utilised to address the current issue, however alternative approaches must be introduced to combat the rise of AMR. Therefore, phytochemicals are another widely used strategy that is just as effective as other antibacterial agents. Previous studies revealed effective antibacterial and anticancer activity by deoxytubulosine which was isolated from Indian medicinal plant *Alangium lamarckii* (Rao and Venkatachalam, 1999). Moreover, corosolic acid also showed anticancer activity with limited side effects (Ma et al., 2018). Our investigation for the phytochemicals which act on *Smlt2267*, *cheW*, *fliM*, *Smlt2318*, and *Smlt2260* revealed that deoxytubulosine and corosolic acid, due to their low binding energy and high affinity, can be used as new antimicrobial agents against resistant strains of *S. maltophilia*.

## CONCLUSION

We have shown protein-protein interaction network comprising 92 virulence genes and 28 AMR genes from *Stenotrophomonas maltophilia K279a* constructed and critically assessed in the current study. Eight hub proteins were identified using comparative topological analysis: *pilL, fliA, Smlt2260, Smlt2267, cheW, Smlt2318, cheZ,* and *fliM*. These proteins will contribute to the discovery of potential therapeutic targets to combat antibiotic resistance. Interestingly, deoxytubulosine and corosolic acid showed better binding affinity towards *Smlt2267, cheW, and fliM,* and *Smlt2318*, and *Smlt2260*, respectively as alternative antibacterial agents for multidrug resistant *S. maltophilia*.

## Acknowledgments and Funding

This work did not receive any specific funding, but RPS gratefully acknowledges the initial funding support from DBT, New Delhi (Grant No: BT/PR41393/MED/30/2298/2020).

## Statement of Ethics

The work is in compliance with ethical standards. No ethical clearance was necessary.

## Data Availability

The sequence data used in this work were obtained from NCBI. The relevant derived data are given in the supplemental tables.

## Author Contributions

SDG, RPS, and RSPR planned the work. LP, RSPR and SDG performed the work and wrote the manuscript. SA helped in data curation. All authors contributed intellectually, and edited/reviewed the manuscript. All authors have read and agreed to the published version of the manuscript.

## Supplemental Information

Supplemental information for this article is available online.

