## Supplemental for "Functional network analysis identifies multiple virulence and antibiotic resistance systems in *Stenotrophomonas maltophilia*"

*Corresponding authors

Running title: Virulence and AMR network in *S. maltophilia*.

***Keywords:*** *Antimicrobial resistance, Functional network analysis, Protein-protein interaction, Virulence*

**Table S1.** AMR genes of *S. maltophilia* K279a.

| Protein ID | CARD AMR  gene name | CARD accession  number | Gene name in GenBank | STRING name |
| --- | --- | --- | --- | --- |
| WP_012481011.1 | aac(6')-Iam | ARO:3002570 | aac(6')-Iz | Smlt3615 |
| WP_012479999.1 | aph(3')-IIc | ARO:3002646 | aph(3')-IIc | Smlt2120 |
| WP_012479999.1 | aph(3')-VIa | ARO:3002652 | aph(3')-IIc | Smlt2120 |
| WP_012480337.1 | aph(6) | ARO:3000151 | Polysaccharide lyase 6 family protein | Smlt2601 |
| WP_012480001.1 | aph(6)-Id | ARO:3002660 | Aminoglycoside phosphotransferase family protein | Smlt2125, spcN |
| WP_005411241.1 | blaA | ARO:3004190 | LysR substrate-binding domain-containing protein | Smlt4229 |
| WP_012480390.1 | blaL1 | ARO:3000582 | blaL1 | Smlt2667 |
| WP_012481073.1 | blaL2 | ARO:3000001 | blaL2, L2 family class A beta-lactamase | Smlt3722 |
| WP_012480939.1 | blaOXA-50 | ARO:3001796 | serine hydrolase | Smlt3495 |
| WP_005410044.1 | ble | ARO:3004256 | Glyoxalase/bleomycin resistance/dioxygenase family protein | Smlt2920 |
| WP_012479057.1 | catB3 | ARO:3002676 | CatB-related O-acetyltransferase | Smlt0620, cat |
| WP_012479057.1 | catB7 | ARO:3002670 | CatB-related O-acetyltransferase | Smlt0620, cat |
| WP_005411585.1 | cmx | ARO:3002703 | MFS transporter | Smlt4538 |
| WP_012479194.1 | dfrA12 | ARO:3002858 | Dihydrofolate reductase | Smlt0814, folA |
| WP_005408768.1 | emrA | ARO:3000027 | emrA | Smlt1529, emrA |
| WP_005408769.1 | emrB | ARO:3000074 | emrB | Smlt1530, emrB |
| WP_012479624.1 | emrC | ARO:3000074 | emrC | Smlt1528 |
| WP_044570606.1 | floR | ARO:3002705 | MFS transporter | Smlt3263 |
| WP_012481544.1 | mexA | ARO:3003051 | smeA | Smlt4476, smeA |
| WP_005409110.1 | mexE | ARO:3000803 | efflux RND transporter periplasmic adaptor subunit | Smlt1830, smeV |
| WP_012481543.1 | mexX | ARO:3003052 | smeB | Smlt4475, smeB |
| WP_012481298.1 | smeF | ARO:3003057 | smeF | Smlt4070, smeF |
| WP_012479732.1 | sul2 | ARO:3003415 | folP | Smlt1734, folP |
| WP_012480512.1 | tet(A) | ARO:3000165 | MFS transporter | Smlt2823 |
| WP_005407901.1 | tet(C) | ARO:3000167 | TetR/AcrR family transcriptional regulator | Smlt0547 |
| WP_012481299.1 | ttgA | ARO:3003055 | smeD | Smlt4072, smeD |
| WP_005411088.1 | ttgB | ARO:3003056 | smeE | Smlt4071, smeE |
| WP_012480988.1 | bcr | ARO:3003801 | multidrug effflux MFS transporter | Smlt3578, bcr |
| WP_099547404.1 | aac(3)-I | ARO:3007384 | aminoglycoside 3-N-acetyltransferase | - |
| WP_004147717.1 | aac(3)-Ia | ARO:3002528 | aminoglycoside 3-N-acetyltransferase | - |
| WP_019237443.1 | aac(2)-Ib | ARO:3002530 | aminoglycoside 3-N-acetyltransferase AAC(2)-Ib | - |
| WP_000557454.1 | aac(3)-IId | ARO:3004623 | aminoglycoside N-acetyltransferase AAC(3)-IId | - |
| QGL76512.1 | aac(3)-IVa | ARO:3002539 | aminoglycoside 3-N-acetyltransferase | - |
| WP_099476052.1 | aac(6')-31 | ARO:3002585 | aminoglycoside N-acetyltransferase AAC(6')-31 | - |
| WP_154331042.1 | aac(6')-Iak | ARO:3003199 | aminoglycoside N-acetyltransferase AAC(6')-Iak | - |
| QGL80238.1 | aac(6')-Ib | ARO:3002546 | aminoglycoside 3-N-acetyltransferase | - |
| QNA95675.1 | aac(6')-Ib' | ARO:3003676 | aminoglycoside 3-N-acetyltransferase | - |
| WP_003159191.1 | aac(6')-Ib4 | ARO:3002577 | aminoglycoside N-acetyltransferase AAC(6')-Ib4 | - |
| UXB37881.1 | aac(6')-Ib-cr5 | ARO:3005115 | aminoglycoside 3-N-acetyltransferase | - |
| AVH90373.1 | aac(6')-Ie/aph(2'')-Ia | ARO:3002597 | aminoglycoside 3-N-acetyltransferase | - |
| WP_023622803.1 | aac(6')-IIa | ARO:3002594 | aminoglycoside N-acetyltransferase AAC(6')-IIa | - |
| QNA95675.1 | aac(6')-IIc | ARO:3002596 | aminoglycoside 3-N-acetyltransferase | - |
| WP_099530169.1 | aac(6')-Iz | ARO:3002570 | aminoglycoside N-acetyltransferase AAC(6')-Iz | - |
| WP_261728213.1 | aadA11 | ARO:3002611 | AadA family aminoglycoside 3''-O-nucleotidyltransferase | - |
| WP_000381802.1 | ant(2'')-Ia | ARO:3000230 | aminoglycoside nucleotidyltransferase ANT(2'')-Ia | - |
| AWB78161.1 | aph(3')-Ia | ARO:3002641 | APH(3')-Ia family aminoglycoside phosphotransferase | - |
| UXF78264.1 | aph(3'')-Ib | ARO:3002639 | APH(3')-Ib family aminoglycoside phosphotransferase | - |
| WP_109815304.1 | aph(3')-IIa | ARO:3002644 | APH(3')-II family aminoglycoside O-phosphotransferase |  |
| QGL92666.1 | aph(3')-IIb | ARO:3002645 | APH(3')-IIb family aminoglycoside O-phosphotransferase | - |
| WP_058199817.1 | aph(3')-VI | ARO:3007412 | APH(3')-VI family aminoglycoside O-phosphotransferase | - |
| WP_069953510.1 | blaAFM-1 | ARO:3006890 | subclass B1 metallo-beta-lactamase AFM-1 | - |
| WP_001931474.1 | blaCARB-2 | ARO:3002241 | PSE family carbenicillin-hydrolyzing class A beta-lactamase CARB-2 | - |
| CAH0065837.1 | blaCTX-M-14 | ARO:3001877 | beta-lactamase | - |
| WP_013250881.1 | blaGES-1 | ARO:3002330 | extended-spectrum class A beta-lactamase GES-1 | - |
| WP_032490683.1 | blaGES-7 | ARO:3002336 | extended-spectrum class A beta-lactamase GES-7 | - |
| WP_000846390.1 | blaOXA-10 | ARO:3001405 | oxacillin-hydrolyzing class D beta-lactamase OXA-10 | - |
| UXY47870.1 | blaOXA-2 | ARO:3001397 | beta-lactamase | - |
| QII30463.1 | blaOXA-35 | ARO:3001429 | beta-lactamase | - |
| UXF78265.1 | blaOXA-46 | ARO:3001797 | beta-lactamase | - |
| WP_063864532.1 | blaOXA-74 | ARO:3001798 | OXA-10 family class D beta-lactamase OXA-74 | - |
| WP_063864599.1 | blaPME-1 | ARO:3001798 | extended-spectrum class A beta-lactamase PME-1 | - |
| AVO31657.1 | blaTEM-116 | ARO:3000979 | broad-spectrum beta-lactamase | - |
| QDY50022.1 | cmlA1 | ARO:3002693 | chloramphenicol efflux MFS transporter CmlA1 | - |
| QDY50022.1 | cmlA10 | - | chloramphenicol efflux MFS transporter CmlA10 | - |
| WP_011977800.1 | cmlA5 | ARO:3002695 | chloramphenicol efflux MFS transporter CmlA5 | - |
| WP_004201280.1 | dfrA14 | ARO:3002859 | trimethoprim-resistant dihydrofolate reductase DfrA14 |  |
| QGL63148.1 | dfrB2 | ARO:3003021 | dihydrofolate reductase | - |
| QNA94854.1 | erm(37) | ARO:3000392 | mrthyltransferase | - |
| WP_024353488.1 | floR2 | ARO:3005043 | chloramphenicol/florfenicol efflux MFS transporter FloR2 | - |
| WP_204311489.1 | fosA | ARO:3000149 | fosfomycin resistance glutathione transferase |  |
| WP_021263608.1 | mph(F) | ARO:3003071 | Mph(F) family macrolide 2'-phosphotransferase | - |
| QQQ40623.1 | msr(E) | ARO:3003109 | ABC-F subfamily protein | - |
| WP_000259031.1 | sul1 | ARO:3000410 | sulfonamide-resistant dihydropteroate synthase Sul1 | - |
| UXF78265.1 | sul3 | ARO:3000413 | sulfonamide-resistant dihydropteroate synthase Sul3 | - |
| WP_230989622.1 | tet(G) | ARO:3000174 | tetracycline efflux MFS transporter Tet(G) | - |

**Table S2.** Virulence genes of *S. maltophilia* K279a.

| Protein ID | VFDB name | Protein name in GenBank | STRING name |
| --- | --- | --- | --- |
| WP_004141526.1 | pilU | pilU | Smlt0612, pilU |
| WP_004147099.1 | pilT | pilT | Smlt1089, pilT |
| WP_004154360.1 | tufA | tuf | Smlt0890, tufB |
| WP_005407653.1 | icl | aceA | Smlt0232, aceA |
| WP_005407764.1 | algC | phosphomannomutase/phosphoglucomutase | Smlt0403, spgM |
| WP_005407915.1 | motA | motA | Smlt0562, motA |
| WP_005408024.1 | xcpR | gspE | Smlt0687, xpsE |
| WP_005408313.1 | acpXL | acpP | Smlt1030, acpP |
| WP_005408982.1 | tviB | Nucleotide sugar dehydrogenase | Smlt1692, wbpO |
| WP_005408999.1 | kdsA | kdsA | Smlt1712, kdsA |
| WP_005409061.1 | panD | Aspartate 1-decarboxylase | Smlt1782, panD |
| WP_005409510.1 | cheB | chemotaxis response regulator protein-glutamate methylesterase | Smlt2248, cheB |
| WP_005409511.1 | cheD | cheD | Smlt2249, cheD |
| WP_005409516.1 | cheW | cheW | Smlt2256, CheW |
| WP_005409527.1 | motC | motC | Smlt2266, MotA |
| WP_005409532.1 | flhG | p loop NTPase | Smlt2271 |
| WP_005409541.1 | fliP | fliP | Smlt2279, fliP |
| WP_005409543.1 | fliN | fliN | Smlt2281, fliN |
| WP_005409544.1 | fliM | fliM | Smlt2282, fliM |
| WP_005409571.1 | flgG | flgG | Smlt2312, flgG |
| WP_005409577.1 | PA3349 | Chemotaxis protein | Smlt2318 |
| WP_005409890.1 | xcpT | GspG | Smlt2746, gspG |
| WP_005410604.1 | mucD | DegQ family serine endoprotease | Smlt3553 |
| WP_005410644.1 | pilH | pilH | Smlt3598 |
| WP_005410715.1 | pilJ | pilJ | Smlt3671, pilJ |
| WP_005410717.1 | pilH | response regulator | Smlt3673, pilH |
| WP_005410795.1 | pilR | sigma-54 dependent transcriptional regulator | Smlt3755, hydG |
| WP_005411088.1 | acrB | smeE | Smlt4071, smeE |
| WP_005411156.1 | tsr | methyl accepting chemotaxis protein | Smlt4138 |
| WP_005411179.1 | sigA/rpoV | rpoD | Smlt4165, rpoD |
| WP_005411226.1 | htpB | groL | Smlt4214, groEL |
| WP_005412452.1 | pilZ | pilZ | Smlt1037, pilZ |
| WP_005413240.1 | tsr | methyl-accepting chemotaxis protein | Smlt2258 |
| WP_005413247.1 | cheY | cheY | Smlt2269, cheY2 |
| WP_005413248.1 | fliA | RNA polymerase sigma factor FliA | Smlt2270, fliA |
| WP_005413257.1 | fliI | FliI/YscN family ATPase | Smlt2286, fliI |
| WP_005413742.1 | cpaF | cpaF | Smlt2873, cpaF |
| WP_005414173.1 | tsr | methyl-accepting chemotaxis protein | Smlt3588 |
| WP_005414416.1 | waaA | waaA | Smlt3930, kdtA |
| WP_005414929.1 | xcpR | GspE/PulE family protein | Smlt4636 |
| WP_005418455.1 | pilG | pilG | Smlt3674, pilG |
| WP_012478733.1 | PA2359 | ntrC | Smlt0159, glnG |
| WP_012478904.1 | mucD | Do family serine endopeptidase | Smlt0381 |
| WP_012479024.1 | motB | motB | Smlt0561, motB |
| WP_012479088.1 | algA | mannose-1-phosphate guanylyltransferase/mannose-6-phosphate isomerase | Smlt0652, xanB |
| WP_012479089.1 | rfbK1 | phosphomannomutase/phosphoglucomutase | Smlt0653, xanA |
| WP_012479289.1 | fabG | fabG | Smlt1029, fabG |
| WP_012479536.1 | katA | catalase | Smlt1385, KatA |
| WP_012479546.1 | tsr | methyl-accepting chemotaxis protein | Smlt1402 |
| WP_012479564.1 | fepA | TonB-dependent siderophore receptor | Smlt1426 |
| WP_012479685.1 | nueA | kdsB | Smlt1642, kdsB |
| WP_012479751.1 | fpvA | TonB-dependent siderophore receptor | Smlt1762 |
| WP_012480090.1 | cheR | chemotaxis protein CheR | Smlt2250, cheR |
| WP_012480091.1 | tsr | methyl-accepting chemotaxis protein | Smlt2251 |
| WP_012480092.1 | tsr | methyl-accepting chemotaxis protein | Smlt2254 |
| WP_012480093.1 | cheA | chemotaxis protein CheA | Smlt2260, cheA |
| WP_012480098.1 | PA1458 | chemotaxis protein CheA | Smlt2267, CheA |
| WP_012480102.1 | flhB | flhB | Smlt2274, flhB |
| WP_012480113.1 | fleQ | sigma-54 dependent transcriptional regulator | Smlt2295 |
| WP_012480117.1 | fliC | flagellin | Smlt2304, fliC |
| WP_012480118.1 | fliC | flagellin | Smlt2305, flaA |
| WP_012480119.1 | fliC | flagellin | Smlt2306 |
| WP_012480126.1 | flgE | flgE | Smlt2314 |
| WP_012480136.1 | clpB | clpB | Smlt2337, clpB |
| WP_012480355.1 | fleQ | sigma-54 dependent transcriptional regulator | Smlt2621 |
| WP_012480358.1 | fapD | C39 family peptidase | Smlt2625 |
| WP_012480379.1 | mgtB | mgtA | Smlt2655, mgtA |
| WP_012480446.1 | exeE | gspE | Smlt2741, gspE |
| WP_012480470.1 | narH | narH | Smlt2773, narH |
| WP_012480471.1 | narG | nitroreductase | Smlt2774, narG |
| WP_012480509.1 | entB | isochorismatase family protein | Smlt2820 |
| WP_012480510.1 | pchD | AMP-binding protein | Smlt2821 |
| WP_012480605.1 | tsr | methyl-accepting chemotaxis protein | Smlt2954, mcpA |
| WP_012481008.1 | PA2359 | prpR | Smlt3611 |
| WP_012481080.1 | PA2371 | clpB | Smlt3732, clpB |
| WP_012481096.1 | pilB | pilB | Smlt3756, pilF |
| WP_012481098.1 | tapA | pilin | Smlt3758, PilE |
| WP_012481099.1 | tapC | type II secretion system F family protein | Smlt3759, PilG |
| WP_012481100.1 | xcpA/pilD | A24 family peptidase | Smlt3760 |
| WP_012481221.1 | flmH | phbB | Smlt3952, PhbB |
| WP_012481316.1 | PA2359 | sigma-54 dependent transcriptional regulator | Smlt4105 |
| WP_012481435.1 | vfr | crp | Smlt4306, cap |
| WP_012481543.1 | acrB | smeB | Smlt4475, smeB |
| WP_012481583.1 | tsr | methyl-accepting chemotaxis protein | Smlt4536 |
| WP_012481655.1 | tsr | methyl-accepting chemotaxis protein | Smlt4645 |
| WP_024958226.1 | fliG | fliG | Smlt2288, fliG |
| WP_043033789.1 | pilU | PilT/PilU family type 4a pilus ATPase | Smlt1090, PilU |
| WP_044570081.1 | flgI | flgI | Smlt2310, flgI |
| WP_044570864.1 | fptA | fhuE | Smlt3999, fhuE |
| WP_044570947.1 | tsr | methyl-accepting chemotaxis protein | Smlt4222 |
| WP_169449112.1 | flhA | flhA | Smlt2273, flhA |
| WP_194434564.1 | pilM | PilM | Smlt3825, pilM |

**Table S3**. *S. maltophilia* AMR genes with their functional partners and confidence scores.

| Target gene | Highest (<0.9 - ≤ 1.0) | High (<0.7 - ≤ 0.9) | Medium (<0.4 - ≤ 0.7) |
| --- | --- | --- | --- |
| Smlt1528 | emrB, emrA | emrB, emrA | smeV, smeB, smeE, smeD, smeA, emrB, emrA |
| emrA | emrB | emrB | emrB |
| folA | folP | folP | folP |
| smeA | smeF, smeE, smeB | smeF, smeE, smeB | smeF, smeE, smeB |
| smeB | smeF, smeD | smeF, smeD, smeV | smeF, smeD, smeV |
| smeD | smeF, smeE | smeF, smeE | smeF, smeE |
| smeE | smeF | smeV, smeF | smeV, smeF |
| Smlt2667 | - | Smlt3722 | Smlt3722 |
| smeF | - | smeV | smeV |
| emrB | - | - | smeB, smeE, smeD, smeA |

**Table S4.** *S. maltophilia* virulence factor genes with their functional partners and confidence scores.

| Target gene | Confidence score | | |
| --- | --- | --- | --- |
|  | **Highest (<0.9 - ≤ 1.0)** | **High (<0.7 - ≤ 0.9)** | **Medium (<0.4 - ≤ 0.7)** |
| Smlt2267 | motB, Smlt1402, cheW, cheB, cheR, Smlt2251, Smlt2254, Smlt2258, Smlt2260, Smlt2266, fliI, flhA, fliM, flhB, fliA, pilJ, Smlt4222, Smlt4536, Smlt4645, Smlt3588, mcpA, Smlt4138, cheY2, Smlt2318 | motB, Smlt0562, Smlt1402, cheW, cheB, cheD, cheR, Smlt2251, Smlt2254, Smlt2258, Smlt2260, Smlt2266, flgG, fliP, Smlt3598, Smlt2314, fliC, Smlt2306, flaA, Smlt2271, Smlt3674, fliG, fliN, fliI, flhA, fliM, flhB, fliA, pilJ, Smlt4222, Smlt4536, Smlt4645, Smlt3588, mcpA, Smlt4138, cheY2, Smlt2318 | motB, Smlt0562, Smlt0612, Smlt1090, Smlt1402, cheW, cheD, cheR, Smlt2251, Smlt2254, Smlt2258, Smlt2260, Smlt2266, pilH, pilM, flgG, fliP, Smlt3598, Smlt2314, fliC, Smlt2306, flaA, Smlt2271, Smlt3674, fliG, fliN, fliI, flhA, fliM, flhB, fliA, pilJ, Smlt4222, Smlt4536, Smlt4645, Smlt3588, cheB, mcpA, Smlt4138, cheY2, Smlt2318 |
| Smlt2266 | motB, fliA, flhB | motB, Smlt0562, Smlt2260, Smlt2314, flgG, Smlt2271, fliC, Smlt2306, flaA, fliI, cheY2, fliP, fliM, fliG, flhA, fliN, fliA, flhB | motB, Smlt0562, cheB, cheR, Smlt2260, flgI, Smlt2318, Smlt2314, flgG, Smlt2271, fliC, Smlt2306, flaA, fliI, cheY2, fliP, fliM, fliG, flhA, fliN, fliA, flhB |
| Smlt3759 | pilT, pilF, Smlt3760 | Smlt0612, xpsE, pilT, Smlt1090, pilE, gspE, pilF, Smlt4636, Smlt3760 | Smlt0612, xpsE, pilT, Smlt1090, pilE, gspE, gspG, pilF, pilM, Smlt4636, Smlt3760 |
| Smlt1402 | Smlt2318, cheD, cheR, cheB, Smlt2260 | cheY2, cheW, Smlt2318, cheD, cheR, cheB, Smlt2260 | pilT, Smlt3598, cheY2, cheW, Smlt2318, cheD, cheR, cheB, Smlt2260 |
| Smlt2251 | cheB, cheD, cheR, Smlt2318, Smlt2260 | cheW, cheB, cheD, cheR, cheY2, Smlt2318, Smlt2260 | motB, cheW, cheB, cheD, cheR, cheY2, Smlt2318, Smlt2260 |
| Smlt2254 | cheB, cheD, cheR, Smlt2318, Smlt2260 | cheW, cheB, cheD, cheR, cheY2, Smlt2318, Smlt2260 | pilT, cheW, cheB, cheD, cheR, fliM, Smlt3598, cheY2, Smlt2318, Smlt2260 |
| Smlt2258 | cheW, cheB, cheD, cheR, Smlt2318, Smlt2260 | cheW, cheB, cheD, cheR, cheY2, Smlt2318, Smlt2260 | cheW, cheB, cheD, cheR, cheY2, Smlt2318, Smlt2260 |
| Smlt2271 | fliA | cheY2, fliA, fliG, Smlt2295, flhB, fliN, fliM, flhA | Smlt2260, cheY2, fliA, fliC, Smlt2306, Smlt2314, Smlt2318, fliI, fliP, fliG, Smlt2295, flhB, fliN, fliM, flhA |
| Smlt2306 | fliA, fliM, flaA | motB, cheB, Smlt2260, fliA, flhA, flhB, fliP, fliN, fliM, fliI, fliG, fliC, flaA, flgG, Smlt2314 | motB, Smlt0562, cheB, cheR, Smlt2260, cheY2, fliA, flhA, fliP, fliN, fliM, fliI, fliG, fliC, flaA, Smlt2318, clpB, flgI, flgG, flhB, Smlt2314 |
| Smlt2314 | fliA, flhA, flhB, fliP, fliN, fliM, fliI, fliG, fliC, flgI, flgG | motB, Smlt0562, Smlt2260, fliA, flhA, flhB, fliP, fliN, fliM, fliI, fliG, fliC, flaA, flgI, flgG, Smlt2318 | motB, Smlt0562, cheR, Smlt2260, cheY2, fliA, flhA, flhB, fliP, fliN, fliM, fliI, fliG, fliC, flaA, flgI, flgG, Smlt2318 |
| Smlt2318 | Smlt2260, Smlt4138 | cheR, Smlt2260, cheY2, fliM, flgI, flgG, Smlt4222, Smlt4536, Smlt3588, Smlt4645, mcpA, Smlt4138 | motB, Smlt0562, cheW, cheB, cheD, cheR, Smlt2260, cheY2, fliA, flhA, flhB, fliP, fliN, fliM, fliI, fliG, fliC, flaA, flgI, flgG, pilJ, Smlt4222, Smlt4536, Smlt3588, Smlt4645, mcpA |
| Smlt2820 | Smlt2821 | Smlt2821 | Smlt4138, Smlt2821 |
| Smlt3588 | cheB, cheD, cheR, Smlt2260 | cheW, cheB, cheD, cheR, Smlt2260, cheY2 | cheW, cheB, cheD, cheR, Smlt2260, cheY2 |
| Smlt3598 | pilJ | Smlt2260, pilH, pilJ | Smlt2260, mcpA, Smlt4222, Smlt4138, Smlt3674, pilH, pilJ |
| Smlt3760 | pilT, pilF | Smlt0612, xpsE, pilT, pilE, gspE, gspG, pilF, Smlt4636 | Smlt0612, xpsE, pilT, pilE, gspE, gspG, pilF, pilM, Smlt4636 |
| Smlt4138 | cheB, cheD, cheR, Smlt2260 | cheW, cheB, cheD, cheR, Smlt2260, cheY2 | pilT, cheW, cheB, cheD, cheR, Smlt2260, cheY2 |
| Smlt4222 | Smlt2260 | cheB, cheD, cheR, Smlt2260 | pilT, cheW, cheB, cheD, cheR, Smlt2260, cheY2 |
| Smlt4536 | cheB, cheD, cheR, Smlt2260 | cheW, cheB, cheD, cheR, Smlt2260, cheY2 | cheW, cheB, cheD, cheR, Smlt2260, cheY2 |
| Smlt4645 | cheB, cheD, cheR, Smlt2260 | cheW, cheB, cheD, cheR, Smlt2260, cheY2 | cheW, cheB, cheD, cheR, Smlt2260, cheY2 |
| acpP | fabG | fabG | fabG |
| cap | rpoD | rpoD | fliA, flhB, pilJ, Smlt3674, rpoD |
| Smlt2260 | motB, cheW, cheB, cheD, cheR, flhB, fliM, fliA, pilJ, cheY2, mcpA | motB, Smlt0562, cheW, cheB, cheD, cheR, Smlt3674, fliC, flaA, fliG, fliN, fliI, flhA, flhB, fliM, fliA, pilJ, cheY2, mcpA | motB, Smlt0562, Smlt0612, cheW, cheB, cheD, cheR, pilH, pilM, flgG, fliP, Smlt3674, fliC, flaA, fliG, fliN, fliI, flhA, flhB, fliM, fliA, pilJ, cheY2, mcpA |
| cheB | mcpA, cheY2, cheD, cheR | motB, cheW, flaA, fliN, fliA, fliM, mcpA, cheY2, cheD, cheR | motB, Smlt0562, cheW, fliI, fliC, fliG, pilJ, flhA, fliP, flhB, flaA, fliN, fliA, fliM, mcpA, cheY2, cheD, cheR |
| cheD | mcpA, cheR | mcpA, cheR | mcpA, cheR |
| cheR | mcpA, cheY2 | motB, Smlt0562, cheW, fliM, flhB, fliN, flhA, fliI, mcpA, cheY2 | motB, Smlt0562, cheW, fliC, flaA, fliA, pilJ, flgG, fliG, fliM, flhB, fliN, flhA, fliI, mcpA, cheY2 |
| cheY2 | fliG, fliA, fliN, fliM | motB, flhB, mcpA, flhA, fliG, fliA, fliN, fliM | motB, Smlt0562, fliP, pilH, flaA, fliC, fliI, flgI, flgG, flhB, mcpA, flhA, fliG, fliA, fliN, fliM |
| clpB | groEL | groEL | flaA, rpoD, groEL |
| flaA | fliA, fliM, fliC | motB, fliA, flhA, flhB, fliP, fliN, fliM, fliI, fliG, fliC, flgG | motB, Smlt0562, fliA, flhA, flhB, fliP, fliN, fliM, fliI, fliG, fliC, flgI, flgG |
| xanA | xanB | xanB | xanB |
| kdsB | kdtA | kdtA | kdtA |
| Smlt0562 | motB | motB | motB |
| narG | narH | narH | narH |
| Smlt3674 | pilJ | pilJ, pilH | pilT, pilJ, pilH, pilM |
| pilH | pilJ | pilJ | Smlt0612, pilT, pilJ |
| spgM | xanA, xanB | xanA, xanB | xanA, xanB |
| flgG | motB, fliA, flhB, flhA, fliP, fliN, fliM, fliI, fliG, fliC, flgI | motB, Smlt0562, fliA, flhA, flhB, fliP, fliN, fliM, fliI, fliG, fliC, flgI | motB, Smlt0562, fliA, flhA, flhB, fliP, fliN, fliM, fliI, fliG, fliC, flgI |
| flgI | motB, fliA, flhB, flhA, fliP, fliN, fliM, fliI, fliG | motB, Smlt0562, fliA, fliC, fliG, fliM, fliN, fliI, fliP, flhB | motB, Smlt0562, fliA, flhA, flhB, fliP, fliN, fliM, fliI, fliG, fliC, pilJ |
| flhA | motB, fliA, fliC, fliG, fliM, fliN, fliI, fliP, flhB |  | motB, Smlt0562, fliA, fliC, fliG, fliM, fliN, fliI, fliP, flhB |
| gspE | gspG | xpsE, gspG | xpsE, pilT, pilM, gspG |
| kdsA | kdsB, kdtA | kdsB, kdtA | xanB, kdsB, kdtA |
| flhB | fliA, fliC, fliN, fliG, fliI, fliM, fliP | motB, Smlt0562, fliA, fliC, fliN, fliG, fliI, fliM, fliP | motB, Smlt0562, fliA, pilJ, fliC, fliN, fliG, fliI, fliM, fliP |
| fliA | fliN, fliC, fliI, fliG, fliM | motB, Smlt0562, fliP, fliN, fliC, fliI, fliG, fliM | motB, Smlt0562, pilJ, fliP, fliN, fliC, fliI, fliG, fliM |
| fliC | fliP, fliN, fliM, fliI, fliG | motB, fliP, fliN, fliM, fliI, fliG | motB, Smlt0562, Smlt0612, fliP, fliN, fliM, fliI, fliG |
| fliG | motB, fliP, fliN, fliM, fliI | motB, Smlt0562, fliP, fliN, fliM, fliI | motB, Smlt0562, fliP, fliN, fliM, fliI |
| fliI | fliP, fliN, fliM | motB, Smlt0562, fliP, fliN, fliM | motB, Smlt0562, fliP, fliN, fliM |
| fliM | motB, fliP, fliN | motB, Smlt0562, fliP, fliN | motB, Smlt0562, fliP, fliN |
| fliN | motB, fliP | motB, Smlt0562, fliP | motB, Smlt0562, fliP |
| Smlt1090 | - | pilT, pilJ, Smlt3760 | Smlt1037, pilT, Smlt4636, Smlt2260, fliC, pilH, pilF, pilM, pilE, pilJ, Smlt3760 |
| Smlt1037 | - | pilM | fabG, acpP, pilH, hydG, pilE, Smlt3598, Smlt3674, pilM |
| Smlt1426 | - | Smlt1762, fhuE, Smlt2820, Smlt2821 | Smlt1762, fhuE, Smlt2820, Smlt2821 |
| Smlt2295 | - | fliA, fliM | fliA, flhA, flhB, fliN, fliM, fliI, fliG, rpoD, cap |
| Smlt4636 | - | xpsE, gspE | Smlt0612, xpsE, pilT, gspE, gspG, pilM |
| cheW | - | flhB, mcpA | pilJ, flhA, fliM, cheY2, fliA, flhB, mcpA |
| clpA | - | groEL | groEL |
| fliP | - | motB | motB, Smlt0562 |
| groEL | - | rpoD | tufB, rpoD |
| pilE | - | pilT, pilM | Smlt0612, pilT, pilF, pilM |
| pilJ | - | Smlt0612, pilT | Smlt0612, pilT, pilM |
| pilM | - | pilT | Smlt0612, xpsE, pilT |
| Smlt0381 | - | - | katA, phbB, fabG, groEL |
| Smlt3553 | - | - | fabG, phbB, groEL |
| gspG | - | - | xpsE, pilF |
| hydG | - | - | pilM |
| pilF | - | - | Smlt0612, pilT, pilM |
| pilT | - | - | Smlt0612, xpsE |
| wbpO | - | - | xanB |

**Table S5.** Top 20 proteins from different parameters of CytoHubba.

| Clustering | Stress | Betweenness | Radiality | Closeness | Eccentricity | Bottleneck | MNC | EPC | Degree | DMNC | MCC |
| --- | --- | --- | --- | --- | --- | --- | --- | --- | --- | --- | --- |
| panD | Smlt2141 | Smlt2141 | pilL | pilL | Smlt2141 | Smlt2141 | pilL | pilL | pilL | flgH | cheZ |
| kdsB | groES | pilL | Smlt2260 | Smlt2260 | Smlt2016 | pilL | Smlt2260 | Smlt2260 | Smlt2260 | flgG | fliG |
| Smlt3611 | emrB | groES | Smlt2267 | Smlt2267 | Smlt4645 | kdsD | Smlt2267 | Smlt2267 | Smlt2267 | flgF | flgD |
| kdtA | pilL | rpoB | Smlt2141 | Smlt2318 | cheW | Smlt2318 | Smlt2318 | Smlt2318 | Smlt2318 | fliL | fliN |
| glnG | Smlt1090 | kdsD | fliA | cheY2 | Smlt0381 | rpoB | cheY2 | fliA | cheY2 | fliQ | flhA |
| Smlt2819 | pilM | fliA | Smlt2318 | fliA | Smlt3588 | clpB | fliA | cheZ | fliA | fliO | fliM |
| Smlt2823 | clpB | xanB | cheY2 | Smlt2141 | Smlt2330 | groES | cheZ | cheY2 | cheZ | fliP | flhB |
| narI | Smlt0612 | entA | CheW | CheW | Smlt2143 | flaA | fliM | fliM | fliM | Smlt0562 | fliA |
| Smlt2621 | Smlt2260 | emrB | cheZ | cheZ | Smlt2295 | CheW | cheR | fliI | cheR | Smlt2309 | fliI |
| narJ | Smlt2267 | Smlt1090 | fliM | fliM | mcpA | entF | CheW | flgC | CheW | Smlt2314 | fliC |
| Smlt4105 | pilT | Smlt0612 | cheR | cheR | cheB | fliA | flhB | flhB | flhB | Smlt2266 | flgC |
| narK | emrA | Smlt2260 | fliI | fliI | Smlt4138 | Smlt0612 | fliG | fliN | fliG | fliF | Smlt2314 |
| Smlt1762 | CheW | Smlt2267 | cheB | cheB | dnaK | xanB | fliN | flgD | fliN | fliH | fliP |
| fhuE | dnaK | entF | flgB | fliC | cheY2 | pilM | fliI | flhF | fliI | flhF | fliR |
| fliQ | fliA | folP | Smlt2306 | flhB | cheR | entA | flhA | cheR | flhA | flgI | flhF |
| flgH | Smlt2306 | dnaK | fliC | flgB | cheZ | folP | cheB | flhA | cheB | flgD | fliF |
| flgF | flaA | narG | flaA | Smlt2306 | Smlt2251 | emrB | fliC | fliF | fliC | MotB | motB |
| fliO | kdsD | clpB | pilJ | fliG | clpP | narG | flgD | motB | flgD | motB | flgG |
| flgG | pilJ | CheW | Smlt2314 | fliN | Smlt2318 | htrB | flgB | flgB | Smlt2141 | fliR | flgB |
| fliL | tolC | pilM | cheW | flhA | Smlt2254 | groEL | fliR | fliG | flgB | flgM | fliL |
| Clustering | Stress | Betweenness | Radiality | Closeness | Eccentricity | Bottleneck | MNC | EPC | Degree | DMNC | MCC |
| panD | Smlt2141 | Smlt2141 | pilL | pilL | Smlt2141 | Smlt2141 | pilL | pilL | pilL | flgH | cheZ |
| kdsB | groES | pilL | Smlt2260 | Smlt2260 | Smlt2016 | pilL | Smlt2260 | Smlt2260 | Smlt2260 | flgG | fliG |
| Smlt3611 | emrB | groES | Smlt2267 | Smlt2267 | Smlt4645 | kdsD | Smlt2267 | Smlt2267 | Smlt2267 | flgF | flgD |
| kdtA | pilL | rpoB | Smlt2141 | Smlt2318 | cheW | Smlt2318 | Smlt2318 | Smlt2318 | Smlt2318 | fliL | fliN |
| glnG | Smlt1090 | kdsD | fliA | cheY2 | Smlt0381 | rpoB | cheY2 | fliA | cheY2 | fliQ | flhA |
| Smlt2819 | pilM | fliA | Smlt2318 | fliA | Smlt3588 | clpB | fliA | cheZ | fliA | fliO | fliM |
| Smlt2823 | clpB | xanB | cheY2 | Smlt2141 | Smlt2330 | groES | cheZ | cheY2 | cheZ | fliP | flhB |

**Table S6.** List of various genes/proteins that contributed to various GO enrichment and pathway enrichment terms*.

| Biological process | Molecular function | Cellular component | KEGG pathway | Pfam protein domains | UniProt keywords | InterPro protein domains |
| --- | --- | --- | --- | --- | --- | --- |
| aac(6')-Iam | **cap** | aac(6')-Iam | **cap** | **cheW** | aac(6')-Iam | **cheB** |
| **aceA** | **cheB** | **aceA** | **cheB** | **clpA** | **acpP** | **cheW** |
| **acpP** | **cheW** | **acpP** | cheD | **clpB** | bcr | **cheY2** |
| bcr | **clpA** | bcr | **cheR** | **glnG** | **cheB** | **clpA** |
| **cap** | **clpB** | **cap** | **cheW** | **gspE** | cheD | **clpB** |
| **cat** | **fabG** | **cat** | **cheY2** | **hydG** | **cheY2** | **flaA** |
| **cheB** | fhuE | **cheB** | **flgG** | mcpA | **clpA** | **fliC** |
| cheD | **fliI** | **cheR** | **flgI** | **pilF** | **clpB** | **fliI** |
| **cheR** | **folA** | **cheW** | **flhA** | **pilJ** | **emrA** | **glnG** |
| **cheW** | **glnG** | **cheY2** | **flhB** | **pilT** | **emrB** | **gspE** |
| **cheY2** | **groEL** | **clpA** | **fliA** | Smlt0612 | **flaA** | **hydG** |
| **clpA** | **gspE** | **clpB** | **fliG** | **Smlt1090** | **flgG** | mcpA |
| **clpB** | **hydG** | **emrA** | **fliI** | **Smlt1402** | **flgI** | **pilF** |
| **emrA** | mcpA | **emrB** | **fliM** | **Smlt2251** | **flhA** | pilG |
| **emrB** | mgtA | **fabG** | **fliN** | **Smlt2254** | **flhB** | **pilH** |
| **fabG** | **motB** | fhuE | fliP | **Smlt2258** | **fliA** | **pilJ** |
| fhuE | **pilF** | **flaA** | **glnG** | **Smlt2260** | **fliC** | **pilT** |
| **flaA** | **pilJ** | **flgG** | **hydG** | **Smlt2267** | **fliG** | Smlt0612 |
| **flgG** | **pilT** | **flgI** | mcpA | **Smlt2295** | **fliM** | **Smlt1090** |
| **flgI** | **Smlt1402** | **flhA** | **motB** | **Smlt2318** | **fliN** | **Smlt1402** |
| **flhB** | **Smlt1762** | **flhB** | **narG** | **Smlt2621** | fliP | **Smlt2251** |
| **fliA** | Smlt2120 | **fliA** | **narH** | Smlt2873 | **glnG** | **Smlt2254** |
| **fliC** | **Smlt2251** | **fliC** | pilG | **Smlt3588** | **gspE** | **Smlt2258** |
| **fliG** | **Smlt2254** | **fliG** | **pilH** | **Smlt3611** | **hydG** | **Smlt2260** |
| **fliI** | **Smlt2258** | **fliI** | **pilJ** | **Smlt4105** | **kdsA** | **Smlt2267** |
| **fliM** | **Smlt2260** | **fliM** | **smeA** | **Smlt4138** | **kdsB** | **Smlt2271** |
| **fliN** | **Smlt2267** | **fliN** | **smeB** | **Smlt4222** | **kdtA** | **Smlt2295** |
| fliP | **Smlt2271** | fliP | **smeD** | **Smlt4536** | mcpA | **Smlt2306** |
| **folA** | **Smlt2295** | **folA** | **smeE** | **Smlt4636** | **pilH** | **Smlt2318** |
| **folP** | **Smlt2318** | **folP** | **smeF** | **Smlt4645** | **pilJ** | **Smlt2621** |
| **glnG** | **Smlt2621** | **glnG** | **Smlt0562** | xpsE | **rpoD** | Smlt2873 |
| **groEL** | Smlt2625 | **groEL** | **Smlt1402** | **-** | **smeA** | **Smlt3588** |
| **gspE** | **Smlt2821** | **gspE** | **Smlt1426** | **-** | **smeB** | **Smlt3598** |
| **gspG** | Smlt2873 | **gspG** | **Smlt2251** | **-** | **smeD** | **Smlt3611** |
| **hydG** | **Smlt3588** | **hydG** | **Smlt2254** | **-** | **smeE** | **Smlt4105** |
| katA | **Smlt3611** | katA | **Smlt2258** | **-** | **smeF** | **Smlt4138** |
| **kdsA** | **Smlt4105** | **kdsA** | **Smlt2260** | **-** | **smeV** | **Smlt4222** |
| **kdsB** | **Smlt4138** | **kdsB** | **Smlt2266** | **-** | **Smlt0562** | **Smlt4536** |
| **kdtA** | **Smlt4222** | **kdtA** | **Smlt2267** | **-** | **Smlt1402** | **Smlt4636** |
| mcpA | **Smlt4536** | mcpA | **Smlt2295** | **-** | **Smlt1426** | **Smlt4645** |
| mgtA | **Smlt4636** | mgtA | **Smlt2314** | **-** | **Smlt1528** | tufB |
| **motB** | **Smlt4645** | **motB** | **Smlt2318** | **-** | **Smlt1762** | xpsE |
| **narG** | spcN | **narG** | **Smlt2601** | **-** | Smlt2120 | **-** |
| **narH** | tufB | **narH** | **Smlt2667** | **-** | **Smlt2251** | **-** |
| panD | **wbpO** | panD | **Smlt3553** | **-** | **Smlt2254** | **-** |
| **pilF** | xanB | phbB | **Smlt3588** | **-** | **Smlt2258** | **-** |
| **pilH** | xpsE | pilE | **Smlt3598** | **-** | **Smlt2260** | **-** |
| **pilJ** | **-** | **pilF** | Smlt3722 | **-** | **Smlt2266** | **-** |
| **pilM** | **-** | **pilH** | **Smlt4138** | **-** | **Smlt2267** | **-** |
| **rpoD** | **-** | **pilJ** | **Smlt4536** | **-** | **Smlt2271** | **-** |
| **smeA** | **-** | **pilT** | **Smlt4645** | **-** | **Smlt2306** | **-** |
| **smeB** | **-** | **rpoD** | **spgM** | **-** | **Smlt2314** | **-** |
| **smeD** | **-** | **smeA** | **xanA** | **-** | **Smlt2318** | **-** |
| **smeE** | **-** | **smeB** | xanB | **-** | **Smlt3588** | **-** |
| **smeF** | **-** | **smeD** | **-** | **-** | **Smlt3598** | **-** |
| **smeV** | **-** | **smeE** | **-** | **-** | **Smlt3674** | **-** |
| Smlt0547 | **-** | **smeF** | **-** | **-** | Smlt3722 | **-** |
| **Smlt0562** | **-** | **smeV** | **-** | **-** | **Smlt3759** | **-** |
| **Smlt1402** | **-** | **Smlt0381** | **-** | **-** | **Smlt4105** | **-** |
| **Smlt1426** | **-** | Smlt0547 | **-** | **-** | **Smlt4138** | **-** |
| **Smlt1528** | **-** | **Smlt0562** | **-** | **-** | **Smlt4222** | **-** |
| **Smlt1762** | **-** | **Smlt1402** | **-** | **-** | Smlt4229 | **-** |
| Smlt2120 | **-** | **Smlt1426** | **-** | **-** | **Smlt4536** | **-** |
| **Smlt2251** | **-** | **Smlt1528** | **-** | **-** | **Smlt4645** | **-** |
| **Smlt2254** | **-** | **Smlt1762** | **-** | **-** | **spgM** | **-** |
| **Smlt2258** | **-** | **Smlt2251** | **-** | **-** | xpsE | **-** |
| **Smlt2260** | **-** | **Smlt2254** | **-** | **-** | **-** | **-** |
| **Smlt2266** | **-** | **Smlt2258** | **-** | **-** | **-** | **-** |
| **Smlt2267** | **-** | **Smlt2260** | **-** | **-** | **-** | **-** |
| **Smlt2271** | **-** | **Smlt2266** | **-** | **-** | **-** | **-** |
| **Smlt2295** | **-** | **Smlt2267** | **-** | **-** | **-** | **-** |
| **Smlt2306** | **-** | **Smlt2271** | **-** | **-** | **-** | **-** |
| **Smlt2314** | **-** | **Smlt2295** | **-** | **-** | **-** | **-** |
| **Smlt2318** | **-** | **Smlt2306** | **-** | **-** | **-** | **-** |
| **Smlt2601** | **-** | **Smlt2314** | **-** | **-** | **-** | **-** |
| **Smlt2621** | **-** | **Smlt2318** | **-** | **-** | **-** | **-** |
| Smlt2625 | **-** | **Smlt2601** | **-** | **-** | **-** | **-** |
| **Smlt2667** | **-** | **Smlt2621** | **-** | **-** | **-** | **-** |
| **Smlt2820** | **-** | Smlt2625 | **-** | **-** | **-** | **-** |
| **Smlt2821** | **-** | **Smlt2667** | **-** | **-** | **-** | **-** |
| **Smlt2823** | **-** | **Smlt2820** | **-** | **-** | **-** | **-** |
| Smlt2873 | **-** | **Smlt2821** | **-** | **-** | **-** | **-** |
| Smlt3263 | **-** | **Smlt2823** | **-** | **-** | **-** | **-** |
| **Smlt3588** | **-** | Smlt2873 | **-** | **-** | **-** | **-** |
| **Smlt3598** | **-** | Smlt2920 | **-** | **-** | **-** | **-** |
| **Smlt3611** | **-** | Smlt3263 | **-** | **-** | **-** | **-** |
| **Smlt3674** | **-** | Smlt3495 | **-** | **-** | **-** | **-** |
| Smlt3722 | **-** | **Smlt3553** | **-** | **-** | **-** | **-** |
| **Smlt3759** | **-** | **Smlt3588** | **-** | **-** | **-** | **-** |
| **Smlt3760** | **-** | **Smlt3598** | **-** | **-** | **-** | **-** |
| **Smlt4105** | **-** | **Smlt3611** | **-** | **-** | **-** | **-** |
| **Smlt4138** | **-** | **Smlt3674** | **-** | **-** | **-** | **-** |
| **Smlt4222** | **-** | Smlt3722 | **-** | **-** | **-** | **-** |
| Smlt4229 | **-** | **Smlt3759** | **-** | **-** | **-** | **-** |
| **Smlt4536** | **-** | **Smlt3760** | **-** | **-** | **-** | **-** |
| Smlt4538 | **-** | **Smlt4105** | **-** | **-** | **-** | **-** |
| **Smlt4636** | **-** | **Smlt4138** | **-** | **-** | **-** | **-** |
| **Smlt4645** | **-** | **Smlt4222** | **-** | **-** | **-** | **-** |
| spcN | **-** | Smlt4229 | **-** | **-** | **-** | **-** |
| **spgM** | **-** | **Smlt4536** | **-** | **-** | **-** | **-** |
| tufB | **-** | Smlt4538 | **-** | **-** | **-** | **-** |
| **wbpO** | **-** | **Smlt4636** | **-** | **-** | **-** | **-** |
| **xanA** | **-** | **Smlt4645** | **-** | **-** | **-** | **-** |
| xanB | **-** | spcN | **-** | **-** | **-** | **-** |
| xpsE | **-** | **spgM** | **-** | **-** | **-** | **-** |
| **-** | **-** | tufB | **-** | **-** | **-** | **-** |
| **-** | **-** | **wbpO** | **-** | **-** | **-** | **-** |
| **-** | **-** | **xanA** | **-** | **-** | **-** | **-** |
| **-** | **-** | xpsE | **-** | **-** | **-** | **-** |

*Target genes in bold.

**Table S7.** Hub proteins functions and subcellular localization.

| Hub protein name | Protein ID | Function | | | | Location | |
| --- | --- | --- | --- | --- | --- | --- | --- |
|  |  | **GenBank** | **Major BLAST hit** | **MOTIF** | **ScanProsite** | **PSLPred** | **PSORTb** |
| pilL | WP_012481044.1 | Hpt domain-containing protein | Histidine kinase | PF01627, Hpt domain | Histidine-containing phosphotransfer (HPt) domain; Histidine kinase domain; Response regulatory domain | Cytoplasmic protein | Cytoplasmic protein |
| fliA | WP_005413248.1 | RNA polymerase sigma factor FliA | RNA polymerase sigma factor for flagellar operon | PF04545, Sigma-70 | Sigma-70 factors family | Cytoplasmic protein | Cytoplasmic protein |
| Smlt2260 | WP_012480093.1 | Chemotaxis protein CheA | Histidine kinase | PF02518, Histidine kinase-, DNA gyrase B-, and HSP90-like ATPase; PF01584, CheW-like domain | Histidine-containing phosphotransfer (HPt) domain; Histidine kinase domain; Phosphopantetheine attachment | Cytoplasmic protein | Cytoplasmic protein |
| Smlt2267 | WP_012480098.1 | Chemotaxis protein CheA | Histidine kinase | PF01584, CheW-like domain | CheW-like domain; Histidine-containing phosphotransfer (HPt) domain; Histidine kinase domain | Cytoplasmic protein | Cytoplasmic protein |
| CheW | WP_005409516.1 | Chemotaxis protein CheW | Putative purine binding chemotaxis protein CheW | PF01584, CheW-like domain | CheW-like domain | Cytoplasmic protein | Cytoplasmic protein |
| Smlt2318 | WP_005409577.1 | Chemotaxis protein | Putative two-component response regulator chemotaxis signal transduction protein | PF01584, CheW-like domain; PF00072, Response regulator receiver domain | Response regulatory domain; CheW-like domain | Cytoplasmic protein | Cytoplasmic protein |
| cheZ | WP_012480099.1 | Protein phosphatase CheZ | Protein phosphatase CheZ | PF04344, Chemotaxis phosphatase, CheZ | no hit | Cytoplasmic protein | Cytoplasmic protein |
| fliM | WP_005409544.1 | Flagellar motor switch protein | Flagellar motor switch protein FliM | PF02154, Flagellar motor switch protein FliM | no hit | Cytoplasmic (This protein may have multiple localization sites) | Cytoplasmic protein |

**Table S8**. Selected ligands for the docking study from medicinal plants described in IMPPAT database.

| Medicinal plant | Selected ligand/phytochemical |
| --- | --- |
| Acalypha indica | 2-Methylanthraquinone |
|  | Asperglaucide |
|  | Kaempferol |
|  | Chrysin |
| Alangium salviifolium | Emetine |
|  | Cephaeline |
|  | Deoxytubulosine |
|  | Amsacrine |
|  | Psychotrine |
|  | Friedelin |
| Berberis aristata | Berberine |
|  | Quercetin |
|  | Jatrorrhizine |
| Calophyllum apetalum | Betulinic acid |
|  | Canophyllol |
| Camellia sinensis | Brassinolide |
|  | 2,2,6-Trimethylcyclohexanone |
|  | Epigallocatechin gallate |
|  | Theasapogenol E |
|  | Theasinensin B |
|  | Theobromine |
|  | Vitexin |
| Cananga odorata | Anonaine |
|  | Viridiflorene |
|  | Viridiflorol |
|  | Eugenol |
|  | Estragole |
| Catunaregam spinosa | Ursolic acid |
| Cinnamomum camphora | Cinnamaldehyde |
|  | Laurolitsine |
| Fagopyrum acutatum | Hecogenin |
| Juniperus indica | Oplopanone |
|  | Abietatriene |
|  | Germacrene B |
|  | Thymol |
|  | Piperitone |
| Mirabilis jalapa | Trigonelline |
| Moringa oleifera | Gossypetin |
|  | Luteolin |
|  | Apigenin |
|  | Quercetagetin |
|  | Niazinin |
| Myrtus communis | Elemicin |
|  | Levomenol |
|  | Catechol |
|  | Corosolic acid |
|  | D-Limonene |
| Ocimum basilicum | Eriodictyol |
|  | Ferulic acid |
|  | Cinnamic acid |
| Oldenlandia umbellata | Anthraquinone |
|  | Purpurin |
| Piper betle | Eremophilene |
|  | Isoeugenol |
|  | Terpinolene |
|  | Aromadendrene |
|  | Diosgenin |
|  | Piperbetol |

**Table S9.** Docking results for hub proteins against select phytochemicals.

| Protein | Rank | Ligand | Binding affinity  (kcal/mol) | Entropy | Energy | | |
| --- | --- | --- | --- | --- | --- | --- | --- |
|  |  |  |  |  | **Inter-molecular** | **Van der Waals** | **Electro-static** |
| Smlt2267 | 1 | Deoxytubulosine | -8.648 | 102.48 | -27.437 | -25.686 | -1.751 |
|  | 2 | Psychotrine | -8.474 | 123.744 | -26.525 | -22.316 | -4.209 |
|  | 3 | Cephaeline | -8.334 | 127.768 | -25.17 | -24.036 | -1.134 |
|  | 4 | Friedelin | -8.301 | 43683.044 | -25.62 | -25.278 | -0.342 |
|  | 5 | Jatrorrhizine | -8.224 | 73.547 | -25.077 | -19.027 | -6.05 |
|  | 6 | Diosgenin | -8.147 | 2705.145 | -26.879 | -19.283 | -7.596 |
|  | 7 | Berberine | -8.134 | 60.932 | -24.201 | -18.338 | -5.863 |
|  | 8 | Abietatriene | -7.97 | 1203.736 | -18.465 | -18.473 | 0.008 |
|  | 9 | Corosolic acid | -7.94 | 969.614 | -30.683 | -15.451 | -15.232 |
|  | 10 | Ursolic acid | -7.907 | 964.232 | -30.66 | -15.459 | -15.201 |
|  | 11 | D-Limonene | -7.824 | 7.169 | -19.459 | -19.444 | -0.015 |
|  | 12 | Estragole | -7.801 | 26.682 | -21.238 | -19.909 | -1.329 |
|  | 13 | Brassinolide | -7.787 | 315.049 | -28.826 | -16.493 | -12.333 |
|  | 14 | Emetine | -7.773 | 142.173 | -25.012 | -19.741 | -5.271 |
|  | 15 | Hecogenin | -7.768 | 16.949 | -23.88 | -17.415 | -6.465 |
|  | 16 | Vitexin | -7.714 | 13232.731 | -31.152 | -15.6 | -15.552 |
|  | 17 | Betulinic acid | -7.712 | 68.999 | -36.286 | -12.371 | -23.915 |
|  | 18 | Theasinensin B | -7.702 | 9799.002 | -33.056 | -10.035 | -23.021 |
|  | 19 | Terpinolene | -7.659 | 132.966 | -39.43 | -10.42 | -29.01 |
|  | 20 | Amsacrine | -7.658 | 60.735 | -17.821 | -16.621 | -1.2 |
|  | 21 | Theasapogenol E | -7.638 | 25.36 | -17.449 | -17.238 | -0.211 |
|  | 22 | Isoeugenol | -7.55 | 77.059 | -30.151 | -4.29 | -25.861 |
|  | 23 | Viridiflorene | -7.545 | 2182.793 | -32.783 | -12.355 | -20.428 |
|  | 24 | Piperitone | -7.539 | 17.198 | -20.077 | -18.567 | -1.51 |
|  | 25 | Canophyllol | -7.487 | 59.228 | -16.108 | -15.894 | -0.214 |
|  | 26 | Levomenol | -7.485 | 12.417 | -18.272 | -18.271 | -0.001 |
|  | 27 | Anonaine | -7.472 | 37645.407 | -25.755 | -11.632 | -14.123 |
|  | 28 | Eremophilene | -7.421 | 36.357 | -20.452 | -10.821 | -9.631 |
|  | 29 | 2,2, 6-Trimethylcyclohexanone | -7.391 | 83.499 | -20.292 | -16.139 | -4.153 |
|  | 30 | Apigenin | -7.343 | 66.786 | -15.377 | -15.186 | -0.191 |
|  | 31 | 2-Methylanthraquinone | -7.339 | 12.264 | -17.236 | -15.864 | -1.372 |
|  | 32 | Cinnamic acid | -7.336 | 18.852 | -26.774 | -11.023 | -15.751 |
|  | 33 | Aromadendrene | -7.316 | 21.625 | -20.526 | -11.996 | -8.53 |
|  | 34 | Germacrene B | -7.3 | -2.818 | -22.058 | -14.124 | -7.934 |
|  | 35 | Piperbetol | -7.299 | 687.694 | -15.505 | -14.74 | -0.765 |
|  | 36 | Viridiflorol | -7.25 | 44348.3 | -17.939 | -16.756 | -1.183 |
|  | 37 | Asperglaucide | -7.233 | 90.055 | -27.018 | -7.718 | -19.3 |
|  | 38 | Anthraquinone | -7.217 | 4360.438 | -15.296 | -15.17 | -0.126 |
|  | 39 | Gossypetin | -7.208 | 96.953 | -24.903 | -13.637 | -11.266 |
|  | 40 | Epigallocatechin gallate | -7.206 | 19.338 | -20.246 | -10.706 | -9.54 |
|  | 41 | Cinnamaldehyde | -7.173 | 29.856 | -35.024 | -7.846 | -27.178 |
|  | 42 | Niazinin | -7.111 | 48.604 | -45.02 | 0.758 | -45.778 |
|  | 43 | Eugenol | -7.107 | 3.621 | -20.6 | -12.748 | -7.852 |
|  | 44 | Ferulic acid | -7.067 | 48.032 | -32.513 | -10.217 | -22.296 |
|  | 45 | Chrysin | -7.027 | 115.103 | -28.265 | -8.306 | -19.959 |
|  | 46 | Quercetagetin | -7.001 | 29.362 | -21.106 | -9.527 | -11.579 |
|  | 47 | Oplopanone | -6.92 | 4.742 | -30.596 | -3.002 | -27.594 |
|  | 48 | Luteolin | -6.781 | -43.557 | -21.173 | -8.842 | -12.331 |
|  | 49 | Kaempferol | -6.754 | 23.577 | -23.808 | -9.817 | -13.991 |
|  | 50 | Quercetin | -6.695 | 26.602 | -31.722 | -10.652 | -21.07 |
|  | 51 | Eriodictyol | -6.641 | 87.547 | -21.504 | -6.732 | -14.772 |
|  | 52 | Catechol | -6.636 | 17.76 | -29.261 | -10.62 | -18.641 |
|  | 53 | Trigonelline | -6.62 | 28.461 | -26.625 | -2.669 | -23.956 |
|  | 54 | Purpurin | -6.586 | 25.064 | -30.924 | -10.602 | -20.322 |
|  | 55 | Elemicin | -6.581 | 24.129 | -30.589 | -11.811 | -18.778 |
|  | 56 | Laurolitsine | -6.49 | -7.917 | -20.319 | -1.152 | -19.167 |
|  | 57 | Theobromine | -6.218 | -14.065 | -28.061 | 4.416 | -32.477 |
|  | 58 | Thymol | -6.186 | 17.892 | -35.075 | -0.382 | -34.693 |
| CheZ | 1 | Cephaeline | -8.137 | 131.537 | -20.983 | -20.173 | -0.81 |
|  | 2 | Psychotrine | -8.008 | 124.548 | -26.129 | -21.745 | -4.384 |
|  | 3 | Abietatriene | -7.973 | 1204.873 | -17.296 | -18.096 | 0.8 |
|  | 4 | Ursolic acid | -7.846 | 967.233 | -30.099 | -17.842 | -12.257 |
|  | 5 | Jatrorrhizine | -7.839 | 70.358 | -31.084 | -11.214 | -19.87 |
|  | 6 | Eremophilene | -7.76 | 67.033 | -15.132 | -13.893 | -1.239 |
|  | 7 | Amsacrine | -7.738 | 78.6 | -28.43 | -14.986 | -13.444 |
|  | 8 | Betulinic acid | -7.691 | 9800.558 | -31.775 | -12.074 | -19.701 |
|  | 9 | Aromadendrene | -7.647 | 689.311 | -13.887 | -13.967 | 0.08 |
|  | 10 | Deoxytubulosine | -7.619 | 108.379 | -21.465 | -18.396 | -3.069 |
|  | 11 | Berberine | -7.607 | 61.729 | -23.395 | -14.231 | -9.164 |
|  | 12 | Theasapogenol E | -7.587 | 2185.889 | -31.052 | -14.975 | -16.077 |
|  | 13 | Emetine | -7.501 | 146.941 | -20.581 | -17.96 | -2.621 |
|  | 14 | Brassinolide | -7.488 | 317.033 | -28.551 | -16.922 | -11.629 |
|  | 15 | Anonaine | -7.458 | 84.376 | -19.414 | -15.868 | -3.546 |
|  | 16 | Friedelin | -7.453 | 43684.17 | -24.494 | -16.009 | -8.485 |
|  | 17 | Corosolic acid | -7.43 | 970.783 | -29.863 | -15.692 | -14.171 |
|  | 18 | Theasinensin B | -7.429 | 131.739 | -46.172 | -8.727 | -37.445 |
|  | 19 | Canophyllol | -7.423 | 37646.786 | -24.384 | -16.011 | -8.373 |
|  | 20 | Laurolitsine | -7.356 | 118.049 | -25.776 | -16.3 | -9.476 |
|  | 21 | Viridiflorol | -7.346 | 4361.823 | -13.911 | -13.976 | 0.065 |
|  | 22 | Estragole | -7.316 | 31.706 | -16.43 | -12.494 | -3.936 |
|  | 23 | 2-Methylanthraquinone | -7.232 | 19.9 | -22.252 | -13.699 | -8.553 |
|  | 24 | Viridiflorene | -7.211 | 61.707 | -13.629 | -13.824 | 0.195 |
|  | 25 | Vitexin | -7.172 | 69.47 | -35.281 | -8.226 | -27.055 |
|  | 26 | Terpinolene | -7.171 | 29.739 | -13.07 | -12.31 | -0.76 |
|  | 27 | Anthraquinone | -7.161 | 18.309 | -21.274 | -12.456 | -8.818 |
|  | 28 | D-Limonene | -7.158 | 13.625 | -13.026 | -11.777 | -1.249 |
|  | 29 | Epigallocatechin gallate | -7.146 | 50.713 | -36.033 | -9.482 | -26.551 |
|  | 30 | Eugenol | -7.125 | 30.188 | -20.164 | -7.782 | -12.382 |
|  | 31 | Elemicin | -7.103 | 60.661 | -17.827 | -13.52 | -4.307 |
|  | 32 | Levomenol | -7.056 | 38.766 | -18.433 | -13.496 | -4.937 |
|  | 33 | Hecogenin | -7.049 | 13237.695 | -26.173 | -15.405 | -10.768 |
|  | 34 | Piperbetol | -7.042 | 88.828 | -27.248 | -15.907 | -11.341 |
|  | 35 | Gossypetin | -6.971 | 30.448 | -27.28 | -7.542 | -19.738 |
|  | 36 | Oplopanone | -6.938 | 87.38 | -21.533 | -10.253 | -11.28 |
|  | 37 | Asperglaucide | -6.901 | 94.627 | -27.997 | -11.21 | -16.787 |
|  | 38 | Isoeugenol | -6.802 | 18.413 | -20.444 | -7.094 | -13.35 |
|  | 39 | Quercetin | -6.774 | 26.487 | -36.121 | 0.902 | -37.023 |
|  | 40 | Niazinin | -6.76 | 54.204 | -25.766 | -7.865 | -17.901 |
|  | 41 | Germacrene B | -6.749 | 44350.885 | -15.354 | -15.035 | -0.319 |
|  | 42 | Chrysin | -6.713 | 20.828 | -28.41 | 0.416 | -28.826 |
|  | 43 | Piperitone | -6.706 | 12.878 | -17.865 | -8.704 | -9.161 |
|  | 44 | Luteolin | -6.704 | 16.97 | -35.832 | 5.539 | -41.371 |
|  | 45 | Apigenin | -6.695 | 18.609 | -27.732 | -1.452 | -26.28 |
|  | 46 | Catechol | -6.672 | -5.46 | -17.878 | -4.613 | -13.265 |
|  | 47 | Diosgenin | -6.662 | 2708.252 | -23.725 | -5.014 | -18.711 |
|  | 48 | Purpurin | -6.649 | 17.888 | -28.412 | -1.132 | -27.28 |
|  | 49 | Quercetagetin | -6.645 | 28.915 | -29.694 | -6.581 | -23.113 |
|  | 50 | Thymol | -6.639 | 22.612 | -18.263 | -4.12 | -14.143 |
|  | 51 | Ferulic acid | -6.588 | 7.639 | -24.436 | -5.591 | -18.845 |
|  | 52 | Eriodictyol | -6.486 | 25.648 | -29.92 | 1.042 | -30.962 |
|  | 53 | 2,2,6-Trimethylcyclohexanone | -6.477 | 13.422 | -16.078 | -8.141 | -7.937 |
|  | 54 | Cinnamic acid | -6.46 | 0.135 | -18.994 | -4.611 | -14.383 |
|  | 55 | Kaempferol | -6.455 | 28.425 | -28.239 | 0.304 | -28.543 |
|  | 56 | Theobromine | -6.27 | -43.186 | -20.802 | -6.88 | -13.922 |
|  | 57 | Cinnamaldehyde | -6.245 | 7.336 | -16.8 | -5.401 | -11.399 |
|  | 58 | Trigonelline | -5.982 | -16.944 | -30.215 | 10.223 | -40.438 |
| CheW | 1 | Deoxytubulosine | -8.793 | 104.572 | -25.965 | -23.647 | -2.318 |
|  | 2 | Cephaeline | -8.748 | 124.683 | -28.715 | -24.267 | -4.448 |
|  | 3 | Friedelin | -8.699 | 43685.214 | -23.45 | -23.322 | -0.128 |
|  | 4 | Emetine | -8.656 | 140.931 | -26.346 | -18.796 | -7.55 |
|  | 5 | Betulinic acid | -8.467 | 9798.353 | -32.898 | -12.26 | -20.638 |
|  | 6 | Theasinensin B | -8.466 | 124.69 | -46.002 | -18.895 | -27.107 |
|  | 7 | Diosgenin | -8.404 | 2705.864 | -26.098 | -19.65 | -6.448 |
|  | 8 | Abietatriene | -8.335 | 1200.698 | -21.478 | -21.004 | -0.474 |
|  | 9 | Theasapogenol E | -8.171 | 2181.271 | -34.127 | -19.457 | -14.67 |
|  | 10 | Canophyllol | -8.144 | 37643.143 | -28.121 | -19.599 | -8.522 |
|  | 11 | Brassinolide | -8.14 | 309.392 | -37.389 | -16.675 | -20.714 |
|  | 12 | Germacrene B | -8.135 | 44348.31 | -17.93 | -17.611 | -0.319 |
|  | 13 | Hecogenin | -8.069 | 13236.593 | -27.028 | -20.547 | -6.481 |
|  | 14 | Terpinolene | -8.027 | 27.348 | -15.462 | -15.379 | -0.083 |
|  | 15 | Estragole | -7.968 | 31.282 | -16.616 | -16.077 | -0.539 |
|  | 16 | Asperglaucide | -7.942 | 94.43 | -27.88 | -21.88 | -6 |
|  | 17 | Berberine | -7.94 | 54.978 | -30.359 | -18.838 | -11.521 |
|  | 18 | Viridiflorol | -7.91 | 4358.497 | -17.237 | -17.316 | 0.079 |
|  | 19 | Aromadendrene | -7.859 | 685.085 | -18.113 | -17.577 | -0.536 |
|  | 20 | D-Limonene | -7.85 | 11.935 | -14.701 | -14.537 | -0.164 |
|  | 21 | Psychotrine | -7.792 | 121.972 | -30.272 | -16.187 | -14.085 |
|  | 22 | Anonaine | -7.728 | 84.045 | -19.746 | -17.741 | -2.005 |
|  | 23 | Epigallocatechin gallate | -7.714 | 44.267 | -51.521 | -13.792 | -37.729 |
|  | 24 | Vitexin | -7.699 | 64.102 | -40.077 | -11.423 | -28.654 |
|  | 25 | Viridiflorene | -7.642 | 59.023 | -16.313 | -15.808 | -0.505 |
|  | 26 | Jatrorrhizine | -7.614 | 65.852 | -35.271 | -12.49 | -22.781 |
|  | 27 | Corosolic acid | -7.612 | 966.025 | -35.895 | -9.408 | -26.487 |
|  | 28 | Luteolin | -7.58 | 11.688 | -36.4 | -12.423 | -23.977 |
|  | 29 | Apigenin | -7.573 | 13.178 | -34.269 | -12.648 | -21.621 |
|  | 30 | Quercetagetin | -7.548 | 23.373 | -43.843 | -11.974 | -31.869 |
|  | 31 | Ursolic acid | -7.543 | 964.711 | -29.785 | -11.183 | -18.602 |
|  | 32 | Kaempferol | -7.532 | 21.929 | -34.455 | -14.524 | -19.931 |
|  | 33 | 2-Methylanthraquinone | -7.516 | 22.766 | -19.385 | -13.152 | -6.233 |
|  | 34 | Eriodictyol | -7.506 | 19.096 | -37.242 | -10.374 | -26.868 |
|  | 35 | Gossypetin | -7.505 | 23.3 | -35.785 | -14.006 | -21.779 |
|  | 36 | Chrysin | -7.488 | 17.342 | -31.804 | -11.156 | -20.648 |
|  | 37 | Quercetin | -7.484 | 20.357 | -36.629 | -13.011 | -23.618 |
|  | 38 | Eremophilene | -7.465 | 66.11 | -16.053 | -15.658 | -0.395 |
|  | 39 | Elemicin | -7.432 | 60.599 | -17.648 | -14.021 | -3.627 |
|  | 40 | Piperbetol | -7.372 | 88.993 | -29.828 | -15.166 | -14.662 |
|  | 41 | Laurolitsine | -7.355 | 112.266 | -33.051 | -5.936 | -27.115 |
|  | 42 | Anthraquinone | -7.291 | 21.779 | -17.805 | -9.436 | -8.369 |
|  | 43 | Eugenol | -7.24 | 28.402 | -20.33 | -13.397 | -6.933 |
|  | 44 | Purpurin | -7.18 | 18.356 | -33.932 | -5.526 | -28.406 |
|  | 45 | Levomenol | -7.179 | 35.316 | -21.616 | -14.133 | -7.483 |
|  | 46 | Oplopanone | -7.136 | 87.167 | -21.773 | -13.203 | -8.57 |
|  | 47 | Cinnamaldehyde | -6.953 | 4.648 | -19.39 | -7.917 | -11.473 |
|  | 48 | Amsacrine | -6.949 | 77.612 | -28.822 | -7.978 | -20.844 |
|  | 49 | Niazinin | -6.903 | 47.388 | -34.737 | -8.859 | -25.878 |
|  | 50 | Thymol | -6.871 | 21.058 | -19.726 | -6.134 | -13.592 |
|  | 51 | Piperitone | -6.851 | 15.361 | -15.33 | -6.538 | -8.792 |
|  | 52 | Isoeugenol | -6.756 | 17.496 | -21.529 | -7.316 | -14.213 |
|  | 53 | 2,2,6-Trimethylcyclohexanone | -6.749 | 14.751 | -14.749 | -4.758 | -9.991 |
|  | 54 | Ferulic acid | -6.72 | -0.435 | -38.228 | -3.289 | -34.939 |
|  | 55 | Cinnamic acid | -6.551 | -4.966 | -29.138 | -3.164 | -25.974 |
|  | 56 | Catechol | -6.209 | -8.405 | -20.762 | 0.122 | -20.884 |
|  | 57 | Theobromine | -6.184 | -43.18 | -20.796 | -6.675 | -14.121 |
|  | 58 | Trigonelline | -6.044 | -16.928 | -30.206 | 10.249 | -40.455 |
| Smlt2318 | 1 | Corosolic acid | -8.833 | 965.441 | -38.403 | -14.473 | -23.93 |
|  | 2 | Ursolic acid | -8.809 | 964.74 | -31.772 | -16.13 | -15.642 |
|  | 3 | Emetine | -8.778 | 139.421 | -29.162 | -23.002 | -6.16 |
|  | 4 | Cephaeline | -8.676 | 122.963 | -31.663 | -24.65 | -7.013 |
|  | 5 | Berberine | -8.478 | 55.174 | -29.887 | -20.804 | -9.083 |
|  | 6 | Brassinolide | -8.385 | 312.211 | -33.18 | -14.95 | -18.23 |
|  | 7 | Diosgenin | -8.279 | 2704.699 | -27.842 | -16.377 | -11.465 |
|  | 8 | Deoxytubulosine | -8.26 | 103.713 | -26.191 | -22.049 | -4.142 |
|  | 9 | Friedelin | -8.204 | 43683.576 | -25.088 | -19.701 | -5.387 |
|  | 10 | Jatrorrhizine | -8.147 | 67.066 | -32.8 | -15.261 | -17.539 |
|  | 11 | Anonaine | -8.145 | 83.52 | -20.27 | -18.766 | -1.504 |
|  | 12 | Abietatriene | -8.109 | 1202.822 | -19.353 | -19.377 | 0.024 |
|  | 13 | Theasapogenol E | -8.036 | 2176.614 | -38.738 | -14.527 | -24.211 |
|  | 14 | Viridiflorene | -7.949 | 58.958 | -16.378 | -16.248 | -0.13 |
|  | 15 | Canophyllol | -7.929 | 37645.728 | -25.523 | -19.21 | -6.313 |
|  | 16 | Elemicin | -7.927 | 59.907 | -18.439 | -17.059 | -1.38 |
|  | 17 | Betulinic acid | -7.898 | 9800.522 | -31.068 | -15.256 | -15.812 |
|  | 18 | Laurolitsine | -7.885 | 115.438 | -27.905 | -14.696 | -13.209 |
|  | 19 | Epigallocatechin gallate | -7.845 | 46.866 | -48.266 | -14.548 | -33.718 |
|  | 20 | Aromadendrene | -7.75 | 686.205 | -16.993 | -16.711 | -0.282 |
|  | 21 | Eremophilene | -7.718 | 65.936 | -16.23 | -16.165 | -0.065 |
|  | 22 | Germacrene B | -7.653 | 44347.735 | -18.504 | -18.426 | -0.078 |
|  | 23 | Terpinolene | -7.62 | 29.246 | -13.563 | -13.858 | 0.295 |
|  | 24 | Psychotrine | -7.615 | 119.758 | -33.201 | -16.218 | -16.983 |
|  | 25 | Luteolin | -7.599 | 12.525 | -36.036 | -11.878 | -24.158 |
|  | 26 | Hecogenin | -7.599 | 13237.918 | -25.867 | -11.666 | -14.201 |
|  | 27 | Eriodictyol | -7.567 | 19.48 | -36.437 | -9.899 | -26.538 |
|  | 28 | Viridiflorol | -7.566 | 4358.383 | -17.351 | -16.424 | -0.927 |
|  | 29 | Chrysin | -7.545 | 18.05 | -31.314 | -10.069 | -21.245 |
|  | 30 | Quercetin | -7.538 | 21.728 | -36.08 | -11.73 | -24.35 |
|  | 31 | Oplopanone | -7.493 | 86.14 | -22.878 | -10.653 | -12.225 |
|  | 32 | Kaempferol | -7.466 | 23.199 | -31.948 | -12.329 | -19.619 |
|  | 33 | Quercetagetin | -7.459 | 23.516 | -44.806 | -6.473 | -38.333 |
|  | 34 | Apigenin | -7.437 | 14.324 | -31.457 | -10.99 | -20.467 |
|  | 35 | Gossypetin | -7.409 | 24.513 | -40.32 | -5.842 | -34.478 |
|  | 36 | Asperglaucide | -7.353 | 93.903 | -29.078 | -16.461 | -12.617 |
|  | 37 | Vitexin | -7.352 | 67.939 | -37.407 | -14.619 | -22.788 |
|  | 38 | Amsacrine | -7.326 | 77.406 | -29.729 | -10.589 | -19.14 |
|  | 39 | Levomenol | -7.293 | 34.98 | -22.204 | -15.347 | -6.857 |
|  | 40 | D-Limonene | -7.271 | 13.9 | -12.73 | -12.238 | -0.492 |
|  | 41 | Niazinin | -7.266 | 46.579 | -33.995 | -8.215 | -25.78 |
|  | 42 | Isoeugenol | -7.265 | 15.038 | -23.892 | -8.488 | -15.404 |
|  | 43 | 2-Methylanthraquinone | -7.238 | 23.212 | -18.939 | -9.478 | -9.461 |
|  | 44 | Anthraquinone | -7.169 | 19.6 | -19.984 | -11.389 | -8.595 |
|  | 45 | Eugenol | -7.079 | 26.655 | -23.897 | -6.195 | -17.702 |
|  | 46 | Piperbetol | -7.059 | 89.603 | -28.148 | -5.713 | -22.435 |
|  | 47 | Purpurin | -6.935 | 16.501 | -27.953 | 2.4 | -30.353 |
|  | 48 | Ferulic acid | -6.867 | 7.096 | -25.064 | -2.004 | -23.06 |
|  | 49 | Estragole | -6.856 | 32.598 | -15.48 | -11.384 | -4.096 |
|  | 50 | Thymol | -6.853 | 20.198 | -20.548 | -5.318 | -15.23 |
|  | 51 | 2,2,6-Trimethylcyclohexanone | -6.712 | 12.589 | -16.911 | -8.615 | -8.296 |
|  | 52 | Piperitone | -6.626 | 13.3 | -17.389 | -9.157 | -8.232 |
|  | 53 | Theasinensin B | -6.619 | 124.838 | -44.237 | -2.199 | -42.038 |
|  | 54 | Catechol | -6.606 | -7.64 | -26.981 | 1.725 | -28.706 |
|  | 55 | Cinnamaldehyde | -6.588 | 7.145 | -16.996 | -7.627 | -9.369 |
|  | 56 | Trigonelline | -6.437 | -18.118 | -31.394 | -2.522 | -28.872 |
|  | 57 | Theobromine | -6.301 | -44.677 | -22.293 | -1.467 | -20.826 |
|  | 58 | Cinnamic acid | -6.209 | -2.981 | -22.021 | -0.256 | -21.765 |
| FliA | 1 | Emetine | -8.796 | 141.687 | -25.53 | -23.604 | -1.926 |
|  | 2 | Psychotrine | -8.63 | 124.137 | -27.044 | -20.964 | -6.08 |
|  | 3 | Cephaeline | -8.465 | 127.094 | -25.535 | -22.319 | -3.216 |
|  | 4 | Anonaine | -8.37 | 81.004 | -22.787 | -20.751 | -2.036 |
|  | 5 | Viridiflorene | -8.162 | 57.056 | -18.28 | -18.129 | -0.151 |
|  | 6 | Deoxytubulosine | -8.065 | 105.036 | -25.335 | -19.994 | -5.341 |
|  | 7 | Corosolic acid | -7.947 | 969.569 | -32.386 | -16.639 | -15.747 |
|  | 8 | Abietatriene | -7.936 | 1201.115 | -21.054 | -19.165 | -1.889 |
|  | 9 | Laurolitsine | -7.906 | 117.313 | -26.148 | -14.723 | -11.425 |
|  | 10 | Canophyllol | -7.834 | 37643.397 | -27.762 | -16.467 | -11.295 |
|  | 11 | Eremophilene | -7.798 | 65.203 | -16.963 | -16.851 | -0.112 |
|  | 12 | Ursolic acid | -7.751 | 965.452 | -30.36 | -10.167 | -20.193 |
|  | 13 | Betulinic acid | -7.695 | 9801.304 | -31.496 | -9.132 | -22.364 |
|  | 14 | Elemicin | -7.684 | 59.701 | -18.616 | -17.884 | -0.732 |
|  | 15 | Theasinensin B | -7.674 | 124.616 | -51.644 | -8.094 | -43.55 |
|  | 16 | Hecogenin | -7.668 | 13233.961 | -29.977 | -18.116 | -11.861 |
|  | 17 | Friedelin | -7.633 | 43682.781 | -25.883 | -14.363 | -11.52 |
|  | 18 | Estragole | -7.623 | 30.2 | -18.194 | -15.254 | -2.94 |
|  | 19 | Aromadendrene | -7.53 | 686.995 | -16.203 | -15.516 | -0.687 |
|  | 20 | Oplopanone | -7.52 | 88.211 | -20.826 | -16.764 | -4.062 |
|  | 21 | Amsacrine | -7.503 | 76.693 | -29.7 | -8.927 | -20.773 |
|  | 22 | Germacrene B | -7.478 | 44347.501 | -18.738 | -19.341 | 0.603 |
|  | 23 | Terpinolene | -7.443 | 27.906 | -14.903 | -14.188 | -0.715 |
|  | 24 | Viridiflorol | -7.434 | 4359.262 | -16.472 | -15.814 | -0.658 |
|  | 25 | 2-Methylanthraquinone | -7.395 | 22.752 | -19.399 | -13.121 | -6.278 |
|  | 26 | Brassinolide | -7.349 | 315.89 | -29.096 | -14.105 | -14.991 |
|  | 27 | Epigallocatechin gallate | -7.331 | 47.538 | -38.319 | -5.535 | -32.784 |
|  | 28 | D-Limonene | -7.234 | 12.172 | -14.454 | -13.834 | -0.62 |
|  | 29 | Diosgenin | -7.219 | 2707.194 | -25.15 | -10.571 | -14.579 |
|  | 30 | Purpurin | -7.209 | 15.185 | -30.078 | -8.156 | -21.922 |
|  | 31 | 2,2,6-Trimethylcyclohexanone | -7.197 | 14.072 | -15.428 | -9.199 | -6.229 |
|  | 32 | Jatrorrhizine | -7.184 | 66.613 | -31.987 | -15.219 | -16.768 |
|  | 33 | Piperitone | -7.172 | 11.157 | -19.623 | -10.616 | -9.007 |
|  | 34 | Theasapogenol E | -7.093 | 2180.475 | -34.685 | -4.889 | -29.796 |
|  | 35 | Berberine | -7.075 | 54.838 | -30.238 | -14.177 | -16.061 |
|  | 36 | Asperglaucide | -7.026 | 93.75 | -28.141 | -10.878 | -17.263 |
|  | 37 | Quercetin | -6.932 | 20.17 | -37.379 | -7.288 | -30.091 |
|  | 38 | Eugenol | -6.928 | 27.18 | -23.636 | -4.539 | -19.097 |
|  | 39 | Vitexin | -6.908 | 67.358 | -37.614 | -0.854 | -36.76 |
|  | 40 | Apigenin | -6.899 | 10.587 | -37.728 | -4.104 | -33.624 |
|  | 41 | Kaempferol | -6.887 | 19.726 | -37.221 | -6.898 | -30.323 |
|  | 42 | Levomenol | -6.837 | 35.736 | -21.567 | -14.072 | -7.495 |
|  | 43 | Piperbetol | -6.797 | 90.403 | -25.856 | -9.629 | -16.227 |
|  | 44 | Chrysin | -6.753 | 17.475 | -31.4 | -5.354 | -26.046 |
|  | 45 | Anthraquinone | -6.747 | 18.302 | -21.282 | -10.718 | -10.564 |
|  | 46 | Gossypetin | -6.744 | 21.316 | -38.827 | -4.665 | -34.162 |
|  | 47 | Luteolin | -6.731 | 11.101 | -37.412 | -5.221 | -32.191 |
|  | 48 | Isoeugenol | -6.722 | 18 | -20.886 | -2.286 | -18.6 |
|  | 49 | Niazinin | -6.692 | 50.847 | -29.803 | -9.915 | -19.888 |
|  | 50 | Cinnamaldehyde | -6.579 | 5.598 | -18.435 | -8.375 | -10.06 |
|  | 51 | Eriodictyol | -6.537 | 22.463 | -33.822 | -2.32 | -31.502 |
|  | 52 | Quercetagetin | -6.528 | 25.87 | -39.192 | 3.031 | -42.223 |
|  | 53 | Ferulic acid | -6.511 | 6.446 | -25.343 | -5.447 | -19.896 |
|  | 54 | Thymol | -6.479 | 21.718 | -19.032 | -1.523 | -17.509 |
|  | 55 | Cinnamic acid | -6.421 | -5.417 | -24.581 | -7.351 | -17.23 |
|  | 56 | Theobromine | -6.321 | -43.082 | -20.698 | -1.777 | -18.921 |
|  | 57 | Catechol | -6.283 | -9.795 | -28.176 | 0.335 | -28.511 |
|  | 58 | Trigonelline | -6.002 | -19.128 | -33.504 | 17.853 | -51.357 |
| Smlt2260 | 1 | Corosolic acid | -9.174 | 43677.559 | -31.104 | -30.71 | -0.394 |
|  | 2 | Friedelin | -8.934 | 968.402 | -31.969 | -26.095 | -5.874 |
|  | 3 | Canophyllol | -8.869 | 37639.749 | -31.625 | -29.744 | -1.881 |
|  | 4 | Abietatriene | -8.827 | 1199.307 | -23.019 | -22.84 | -0.179 |
|  | 5 | Betulinic acid | -8.812 | 9800.845 | -30.7 | -23.387 | -7.313 |
|  | 6 | Hecogenin | -8.803 | 13231.058 | -32.729 | -20.426 | -12.303 |
|  | 7 | Ursolic acid | -8.8 | 963.529 | -31.773 | -24.75 | -7.023 |
|  | 8 | Deoxytubulosine | -8.729 | 101.831 | -28.899 | -24.313 | -4.586 |
|  | 9 | Apigenin | -8.623 | 11.39 | -35.709 | -18.083 | -17.626 |
|  | 10 | Cephaeline | -8.467 | 121.654 | -31.013 | -22.938 | -8.075 |
|  | 11 | Emetine | -8.379 | 137.609 | -29.969 | -21.332 | -8.637 |
|  | 12 | Theasapogenol E | -8.368 | 2184.665 | -33.701 | -12.155 | -21.546 |
|  | 13 | 2-Methylanthraquinone | -8.137 | 19.908 | -22.242 | -13.638 | -8.604 |
|  | 14 | Laurolitsine | -8.13 | 114.376 | -28.511 | -14.89 | -13.621 |
|  | 15 | Eremophilene | -8.125 | 64.518 | -17.645 | -17.986 | 0.341 |
|  | 16 | Anthraquinone | -8.056 | 18.146 | -21.438 | -13.424 | -8.014 |
|  | 17 | Levomenol | -7.993 | 35.461 | -21.353 | -20.45 | -0.903 |
|  | 18 | Berberine | -7.917 | 57.16 | -27.909 | -17.937 | -9.972 |
|  | 19 | Germacrene B | -7.853 | 44347.021 | -19.219 | -18.804 | -0.415 |
|  | 20 | Kaempferol | -7.84 | 26.365 | -28.876 | -7.087 | -21.789 |
|  | 21 | Amsacrine | -7.828 | 75.065 | -31.516 | -20.128 | -11.388 |
|  | 22 | Estragole | -7.786 | 30.13 | -18.008 | -16.058 | -1.95 |
|  | 23 | Theasinensin B | -7.762 | 127.445 | -47.874 | -15.606 | -32.268 |
|  | 24 | Viridiflorene | -7.733 | 57.788 | -17.549 | -17.467 | -0.082 |
|  | 25 | Terpinolene | -7.725 | 26.948 | -15.861 | -15.296 | -0.565 |
|  | 26 | Asperglaucide | -7.641 | 96.143 | -25.922 | -20.656 | -5.266 |
|  | 27 | Jatrorrhizine | -7.621 | 64.157 | -36.564 | -15.385 | -21.179 |
|  | 28 | Elemicin | -7.618 | 59.054 | -19.59 | -16.944 | -2.646 |
|  | 29 | Vitexin | -7.588 | 65.777 | -41.909 | -13.917 | -27.992 |
|  | 30 | 2,2,6-Trimethylcyclohexanone | -7.561 | 11.852 | -17.648 | -6.54 | -11.108 |
|  | 31 | Aromadendrene | -7.546 | 685.521 | -17.677 | -16.845 | -0.832 |
|  | 32 | Ferulic acid | -7.541 | 3.667 | -28.212 | -9.912 | -18.3 |
|  | 33 | Brassinolide | -7.52 | 310.559 | -33.772 | -13.596 | -20.176 |
|  | 34 | Piperitone | -7.468 | 10.744 | -19.95 | -9.984 | -9.966 |
|  | 35 | Viridiflorol | -7.444 | 4356.943 | -18.79 | -18.337 | -0.453 |
|  | 36 | D-Limonene | -7.427 | 12.049 | -14.577 | -14.26 | -0.317 |
|  | 37 | Diosgenin | -7.417 | 2702.827 | -29.248 | -16.269 | -12.979 |
|  | 38 | Anonaine | -7.289 | 81.418 | -22.372 | -13.651 | -8.721 |
|  | 39 | Cinnamic acid | -7.252 | -7.782 | -26.83 | -7.898 | -18.932 |
|  | 40 | Psychotrine | -7.096 | 122.556 | -29.611 | -14.293 | -15.318 |
|  | 41 | Catechol | -7.088 | -10.811 | -23.181 | -8.019 | -15.162 |
|  | 42 | Piperbetol | -6.984 | 90.844 | -25.271 | -14.64 | -10.631 |
|  | 43 | Niazinin | -6.948 | 47.282 | -32.855 | -13.148 | -19.707 |
|  | 44 | Quercetagetin | -6.856 | 23.475 | -44.492 | -1.053 | -43.439 |
|  | 45 | Epigallocatechin gallate | -6.788 | 45.947 | -48.529 | -0.625 | -47.904 |
|  | 46 | Gossypetin | -6.781 | 25.792 | -34.468 | -2.288 | -32.18 |
|  | 47 | Chrysin | -6.731 | 19.168 | -30.664 | -5.337 | -25.327 |
|  | 48 | Quercetin | -6.723 | 22.658 | -35.967 | -3.616 | -32.351 |
|  | 49 | Oplopanone | -6.644 | 87.647 | -21.654 | -11.924 | -9.73 |
|  | 50 | Luteolin | -6.623 | 11.298 | -37.621 | -3.756 | -33.865 |
|  | 51 | Purpurin | -6.572 | 16.456 | -35.318 | 1.236 | -36.554 |
|  | 52 | Eriodictyol | -6.568 | 17.876 | -38.893 | -1.821 | -37.072 |
|  | 53 | Isoeugenol | -6.482 | 15.153 | -23.944 | -8.193 | -15.751 |
|  | 54 | Eugenol | -6.451 | 26.924 | -23.821 | -7.926 | -15.895 |
|  | 55 | Thymol | -6.45 | 20.47 | -20.444 | -3.608 | -16.836 |
|  | 56 | Cinnamaldehyde | -6.351 | 5.58 | -18.488 | -7.705 | -10.783 |
|  | 57 | Trigonelline | -6.279 | -17.91 | -31.18 | 4.301 | -35.481 |
|  | 58 | Theobromine | -6.081 | -43.431 | -21.048 | -1.273 | -19.775 |
| FliM | 1 | Deoxytubulosine | -9.579 | 102.305 | -28.387 | -26.248 | -2.139 |
|  | 2 | Emetine | -9.032 | 142.696 | -24.966 | -22.44 | -2.526 |
|  | 3 | Cephaeline | -8.941 | 128.586 | -24.759 | -22.395 | -2.364 |
|  | 4 | Jatrorrhizine | -8.444 | 70.064 | -28.53 | -20.005 | -8.525 |
|  | 5 | Amsacrine | -8.42 | 77.785 | -29.348 | -16.41 | -12.938 |
|  | 6 | Berberine | -8.406 | 57.897 | -27.14 | -18.819 | -8.321 |
|  | 7 | Abietatriene | -8.281 | 1204.429 | -17.796 | -17.429 | -0.367 |
|  | 8 | Epigallocatechin gallate | -8.093 | 47.59 | -40.121 | -14.706 | -25.415 |
|  | 9 | Friedelin | -8.021 | 43686.823 | -21.84 | -19.588 | -2.252 |
|  | 10 | Germacrene B | -7.969 | 44347.928 | -18.312 | -17.734 | -0.578 |
|  | 11 | Laurolitsine | -7.962 | 117.141 | -26.701 | -14.546 | -12.155 |
|  | 12 | Eremophilene | -7.947 | 67.352 | -14.812 | -15.02 | 0.208 |
|  | 13 | Anonaine | -7.925 | 82.123 | -21.668 | -13.4 | -8.268 |
|  | 14 | Elemicin | -7.868 | 60.553 | -18.892 | -14.633 | -4.259 |
|  | 15 | Psychotrine | -7.826 | 122.518 | -29.592 | -12.92 | -16.672 |
|  | 16 | Viridiflorene | -7.715 | 60.586 | -14.75 | -14.371 | -0.379 |
|  | 17 | Terpinolene | -7.641 | 29.304 | -13.505 | -12.84 | -0.665 |
|  | 18 | Estragole | -7.636 | 33.334 | -14.582 | -13.294 | -1.288 |
|  | 19 | Theasinensin B | -7.631 | 133.1 | -37.927 | -11.466 | -26.461 |
|  | 20 | Luteolin | -7.543 | 13.741 | -39.525 | -10.954 | -28.571 |
|  | 21 | Aromadendrene | -7.532 | 689.122 | -14.076 | -13.973 | -0.103 |
|  | 22 | Apigenin | -7.482 | 14.688 | -32.516 | -12.237 | -20.279 |
|  | 23 | Betulinic acid | -7.453 | 9803.175 | -28.315 | -8.883 | -19.432 |
|  | 24 | Chrysin | -7.451 | 20.305 | -28.718 | -9.421 | -19.297 |
|  | 25 | Gossypetin | -7.436 | 25.669 | -39.323 | -9.472 | -29.851 |
|  | 26 | Kaempferol | -7.415 | 24.618 | -31.682 | -12.344 | -19.338 |
|  | 27 | Canophyllol | -7.402 | 37647.874 | -24.202 | -14.118 | -10.084 |
|  | 28 | Quercetin | -7.38 | 23.476 | -39.662 | -7.059 | -32.603 |
|  | 29 | D-Limonene | -7.357 | 14.557 | -12.102 | -12.173 | 0.071 |
|  | 30 | Piperbetol | -7.349 | 86.678 | -30.387 | -11.44 | -18.947 |
|  | 31 | Viridiflorol | -7.313 | 4361.569 | -14.165 | -14.171 | 0.006 |
|  | 32 | Niazinin | -7.264 | 50.765 | -29.587 | -12.146 | -17.441 |
|  | 33 | Diosgenin | -7.261 | 2706.508 | -26.544 | -12.205 | -14.339 |
|  | 34 | 2-Methylanthraquinone | -7.199 | 20.381 | -21.769 | -6.492 | -15.277 |
|  | 35 | Brassinolide | -7.198 | 315.992 | -28.88 | -11.691 | -17.189 |
|  | 36 | Theasapogenol E | -7.174 | 2185.438 | -29.826 | -8.711 | -21.115 |
|  | 37 | Piperitone | -7.125 | 15.023 | -16.925 | -7.205 | -9.72 |
|  | 38 | 2,2,6-Trimethylcyclohexanone | -7.105 | 13.66 | -15.84 | -9.021 | -6.819 |
|  | 39 | Hecogenin | -7.076 | 13236.775 | -26.969 | -11.814 | -15.155 |
|  | 40 | Corosolic acid | -7.056 | 969.823 | -29.856 | -13.087 | -16.769 |
|  | 41 | Isoeugenol | -7.037 | 16.834 | -22.065 | -7.919 | -14.146 |
|  | 42 | Anthraquinone | -7.036 | 18.846 | -20.737 | -6.087 | -14.65 |
|  | 43 | Ursolic acid | -7.001 | 966.504 | -28.136 | -12.477 | -15.659 |
|  | 44 | Thymol | -6.993 | 20.871 | -19.864 | -7.836 | -12.028 |
|  | 45 | Purpurin | -6.972 | 18.064 | -32.273 | -1.849 | -30.424 |
|  | 46 | Levomenol | -6.97 | 38.585 | -18.462 | -11.616 | -6.846 |
|  | 47 | Eugenol | -6.922 | 28.624 | -21.992 | -8.361 | -13.631 |
|  | 48 | Ferulic acid | -6.901 | 7.91 | -25.674 | -8.149 | -17.525 |
|  | 49 | Eriodictyol | -6.9 | 24.763 | -29.839 | -5.868 | -23.971 |
|  | 50 | Cinnamaldehyde | -6.799 | 7.188 | -16.808 | -5.749 | -11.059 |
|  | 51 | Quercetagetin | -6.694 | 24.605 | -40.065 | 5.008 | -45.073 |
|  | 52 | Oplopanone | -6.675 | 87.545 | -21.44 | -3.278 | -18.162 |
|  | 53 | Asperglaucide | -6.514 | 97.848 | -23.953 | -10.924 | -13.029 |
|  | 54 | Vitexin | -6.487 | 69.418 | -35.443 | -0.983 | -34.46 |
|  | 55 | Cinnamic acid | -6.346 | -4.581 | -23.883 | 1.527 | -25.41 |
|  | 56 | Theobromine | -6.211 | -41.704 | -19.32 | -3.042 | -16.278 |
|  | 57 | Catechol | -6.082 | -10.791 | -25.692 | 4.367 | -30.059 |
|  | 58 | Trigonelline | -5.712 | -18.094 | -31.365 | 18.587 | -49.952 |
| PilL | 1 | Emetine | -7.988 | 148.959 | -18.21 | -16.243 | -1.967 |
|  | 2 | Cephaeline | -7.958 | 134.205 | -18.494 | -16.464 | -2.03 |
|  | 3 | Deoxytubulosine | -7.545 | 111.869 | -20.437 | -12.062 | -8.375 |
|  | 4 | Psychotrine | -7.524 | 133.443 | -18.036 | -9.785 | -8.251 |
|  | 5 | Theasinensin B | -7.457 | 144.328 | -24.628 | -14.78 | -9.848 |
|  | 6 | Abietatriene | -7.441 | 1209.741 | -12.423 | -12.125 | -0.298 |
|  | 7 | Berberine | -7.428 | 69.182 | -15.883 | -11.361 | -4.522 |
|  | 8 | Betulinic acid | -7.399 | 9810.799 | -20.83 | -10.625 | -10.205 |
|  | 9 | Laurolitsine | -7.389 | 124.996 | -18.131 | -11.27 | -6.861 |
|  | 10 | Corosolic acid | -7.221 | 981.022 | -18.647 | -8.803 | -9.844 |
|  | 11 | Jatrorrhizine | -7.199 | 82.424 | -16.193 | -9.101 | -7.092 |
|  | 12 | Ursolic acid | -7.155 | 976.771 | -17.733 | -7.654 | -10.079 |
|  | 13 | Quercetagetin | -7.131 | 39.167 | -20.978 | -8.719 | -12.259 |
|  | 14 | Diosgenin | -7.128 | 2714.759 | -17.216 | -9.315 | -7.901 |
|  | 15 | Gossypetin | -7.113 | 39.311 | -18.575 | -8.668 | -9.907 |
|  | 16 | Epigallocatechin gallate | -7.075 | 60.181 | -25.218 | -5.292 | -19.926 |
|  | 17 | Canophyllol | -7.015 | 37654.912 | -16.254 | -10.074 | -6.18 |
|  | 18 | Viridiflorene | -7.003 | 65.323 | -10.013 | -10.155 | 0.142 |
|  | 19 | Eremophilene | -6.955 | 72.478 | -9.688 | -9.927 | 0.239 |
|  | 20 | Anonaine | -6.938 | 87.941 | -15.85 | -6.459 | -9.391 |
|  | 21 | Germacrene B | -6.877 | 44354.731 | -11.509 | -11.713 | 0.204 |
|  | 22 | Brassinolide | -6.873 | 326.232 | -18.006 | -5.101 | -12.905 |
|  | 23 | Eugenol | -6.866 | 35.642 | -13.242 | -8.039 | -5.203 |
|  | 24 | Aromadendrene | -6.851 | 694.468 | -8.73 | -8.489 | -0.241 |
|  | 25 | Niazinin | -6.833 | 60.158 | -18.544 | -8.833 | -9.711 |
|  | 26 | Asperglaucide | -6.825 | 103.736 | -17.813 | -6.872 | -10.941 |
|  | 27 | D-Limonene | -6.81 | 18.668 | -7.959 | -7.848 | -0.111 |
|  | 28 | Piperbetol | -6.798 | 96.032 | -19.487 | -5.323 | -14.164 |
|  | 29 | Vitexin | -6.766 | 82.479 | -20.833 | -5.464 | -15.369 |
|  | 30 | Elemicin | -6.752 | 65.475 | -13.941 | -7.656 | -6.285 |
|  | 31 | Luteolin | -6.72 | 24.983 | -22.054 | -0.585 | -21.469 |
|  | 32 | Amsacrine | -6.718 | 85.595 | -20.729 | -3.761 | -16.968 |
|  | 33 | Kaempferol | -6.702 | 34.114 | -21.585 | -0.079 | -21.506 |
|  | 34 | Quercetin | -6.699 | 34.535 | -21.711 | -0.164 | -21.547 |
|  | 35 | Viridiflorol | -6.69 | 4366.471 | -9.263 | -8.933 | -0.33 |
|  | 36 | Chrysin | -6.683 | 29.593 | -19.053 | -3.558 | -15.495 |
|  | 37 | Apigenin | -6.649 | 24.466 | -21.623 | 0.483 | -22.106 |
|  | 38 | Hecogenin | -6.629 | 13243.843 | -20.177 | -5.938 | -14.239 |
|  | 39 | Eriodictyol | -6.627 | 33.08 | -21.7 | -0.018 | -21.682 |
|  | 40 | Oplopanone | -6.621 | 92.658 | -16.305 | -4.158 | -12.147 |
|  | 41 | Purpurin | -6.615 | 25.062 | -19.319 | -2.943 | -16.376 |
|  | 42 | 2-Methylanthraquinone | -6.615 | 26.125 | -16.026 | -3.667 | -12.359 |
|  | 43 | Friedelin | -6.592 | 43693.194 | -15.47 | -3.139 | -12.331 |
|  | 44 | Anthraquinone | -6.563 | 23.78 | -15.804 | -3.428 | -12.376 |
|  | 45 | Levomenol | -6.534 | 42.14 | -14.645 | -5.768 | -8.877 |
|  | 46 | Ferulic acid | -6.505 | 17.605 | -14.305 | -5.715 | -8.59 |
|  | 47 | Estragole | -6.467 | 37.037 | -10.91 | -6.458 | -4.452 |
|  | 48 | Terpinolene | -6.379 | 34.086 | -8.723 | -6.396 | -2.327 |
|  | 49 | Theasapogenol E | -6.361 | 2197.008 | -20.077 | -4.06 | -16.017 |
|  | 50 | Isoeugenol | -6.353 | 24.687 | -14.23 | -2.563 | -11.667 |
|  | 51 | Piperitone | -6.327 | 16.215 | -14.481 | -1.572 | -12.909 |
|  | 52 | Cinnamaldehyde | -6.303 | 10.299 | -13.72 | -0.577 | -13.143 |
|  | 53 | Thymol | -6.286 | 27.595 | -13.599 | -1.279 | -12.32 |
|  | 54 | Trigonelline | -6.281 | -13.227 | -26.511 | 1.26 | -27.771 |
|  | 55 | Theobromine | -6.231 | -38.538 | -16.154 | -2.87 | -13.284 |
|  | 56 | 2,2,6-Trimethylcyclohexanone | -6.225 | 15.151 | -14.349 | -2.016 | -12.333 |
|  | 57 | Cinnamic acid | -5.916 | 5.174 | -13.845 | 0.248 | -14.093 |
|  | 58 | Catechol | -5.723 | -0.951 | -20.533 | 6.85 | -27.383 |

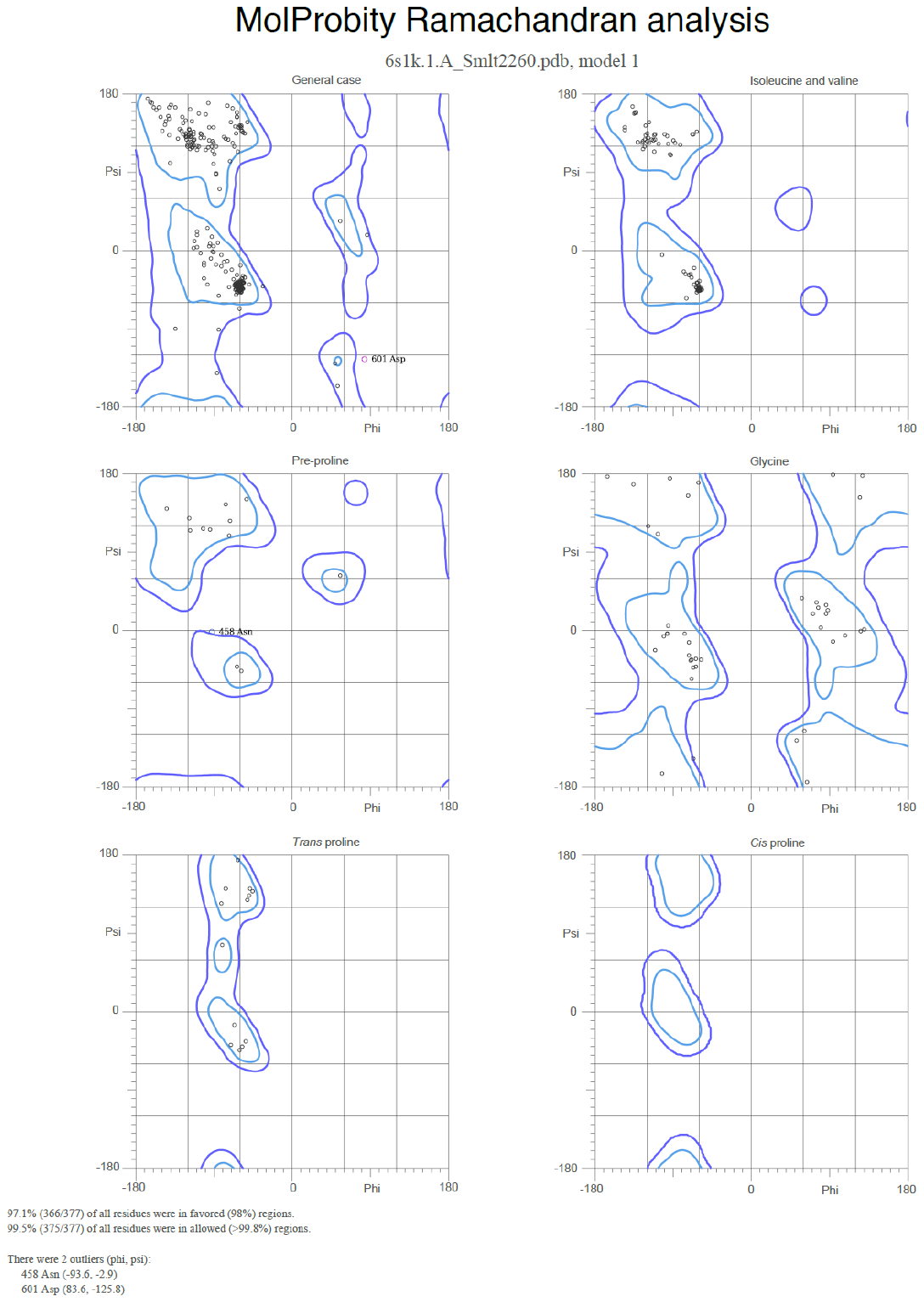

**Fig. S1.** Ramachandran plot of Smlt2260 showing the highest percentage of favored regions as predicted using MOLPROBITY.
